# Neurotransmitter regulation rather than cell-intrinsic properties shape the high-pass filtering properties of olfactory bulb glomeruli

**DOI:** 10.1101/2021.09.14.460371

**Authors:** Joseph D. Zak, Nathan E. Schoppa

## Abstract

GABAergic periglomerular (PG) cells in the olfactory bulb are proposed to mediate an intraglomerular “high-pass” filter through inhibition targeted onto a glomerulus. With this mechanism, weak stimuli (e.g., an odor with a low affinity for an odorant receptor) mainly produce PG cell-driven inhibition but strong stimuli generate enough excitation to overcome inhibition. PG cells may be particularly susceptible to being activated by weak stimuli due to their intrinsically small size and high input resistance. Here, we used dual-cell patch-clamp recordings and imaging methods in bulb slices obtained from wild-type and transgenic rats with labeled GABAergic cells to test a number of predictions of the high-pass filtering model. We first directly compared the responsiveness of PG cells with that of external tufted cells (eTCs), which are a class of excitatory interneurons that reside in a parallel but opposing position in the glomerular circuitry. PG cells were in fact found to be no more responsive than eTCs at low levels of sensory neuron activity. While PG cells required smaller currents to be excited, this advantage was offset by the fact that a given level of sensory neuron activity produced much smaller synaptic currents. We did however identify other factors that shaped the excitation/inhibition balance in a manner that would support a high-pass filter, including glial glutamate transporters and presynaptic metabotropic glutamate receptors. We conclude that a single glomerulus may act as a high-pass filter to enhance the contrast between different olfactory stimuli through mechanisms that are largely independent cell intrinsic properties.

**Key Points Summary:**

- GABAergic periglomerular (PG) cells in the olfactory bulb are proposed to mediate a “high-pass” filter at a single glomerulus that selectively blocks weak stimulus signals. Their efficacy may reflect their intrinsically small size and high input resistance, which allows them to be easily excited.
- We found that PG cells were in fact no more likely to be activated by weak stimuli than excitatory neurons. PG cells spiked more readily in response to a fixed current input, but this advantage for excitability was offset by small synaptic currents.
- Glomeruli nevertheless display an excitation/inhibition balance that can support a high pass filter, becoming much more favorable with increasing stimulus strength. Factors shaping the filter include glial glutamate transporters and presynaptic metabotropic glutamate receptors.
- We conclude that a single glomerulus may act as a high-pass filter to enhance stimulus contrast through mechanisms that are largely independent of cell-intrinsic properties.

## Introduction

Within the mammalian olfactory bulb, GABAergic periglomerular (PG) cells constitute a class of inhibitory interneurons in the glomerular layer that are noted by their axon-less morphology and dendritic arborization that is most often confined to just one glomerulus (Pinching and Powell, 1971; Shepherd et al., 2004; Kiyokage et al., 2010). Functional studies have begun to implicate PG cells as having an important role in olfactory information processing. For example, optogenetic silencing of PG cells greatly reduces odor-evoked inhibitory responses in output mitral cells (MCs) and tufted cells (TCs; Fukunaga et al., 2014) that are widely believed to underlie olfactory contrast enhancement (Yokoi et al., 1995). Computational studies that have modeled a single glomerulus (Cleland and Sethupathy, 2006; Cleland and Linster, 2012) have suggested a more defined function for PG cells that is based on an odor’s affinity for the odorant receptor (OR) encoded by a glomerulus. In this model, odors with a low affinity for the OR mainly excite PG cells, resulting in mainly inhibition and block of an output signal, but higher affinity odors generate enough MC/TC excitation to overcome inhibition (**Fig 1A**). Hence, through modifications in the intraglomerular excitation/inhibition (E/I) balance, a single glomerulus effectively acts as a high-pass filter that selectively enables passage of signals generated by strong stimuli. It has been suggested (Cleland and Sethupathy, 2006; Gire and Schoppa, 2009; Fukunaga et al., 2014) that selective activation of PG cells by weak stimuli is achieved due to their very small size and high input resistance (0.5-2 GΩ; Puopolo and Belluzzi, 1998; McQuiston and Katz, 2001; Smith and Jahr, 2002; Hayar et al., 2004b). These properties might enable PG cells to be more easily excited at weak levels of olfactory sensory neuron (OSN) activity than excitatory cells in the bulb.

**Figure 1.**
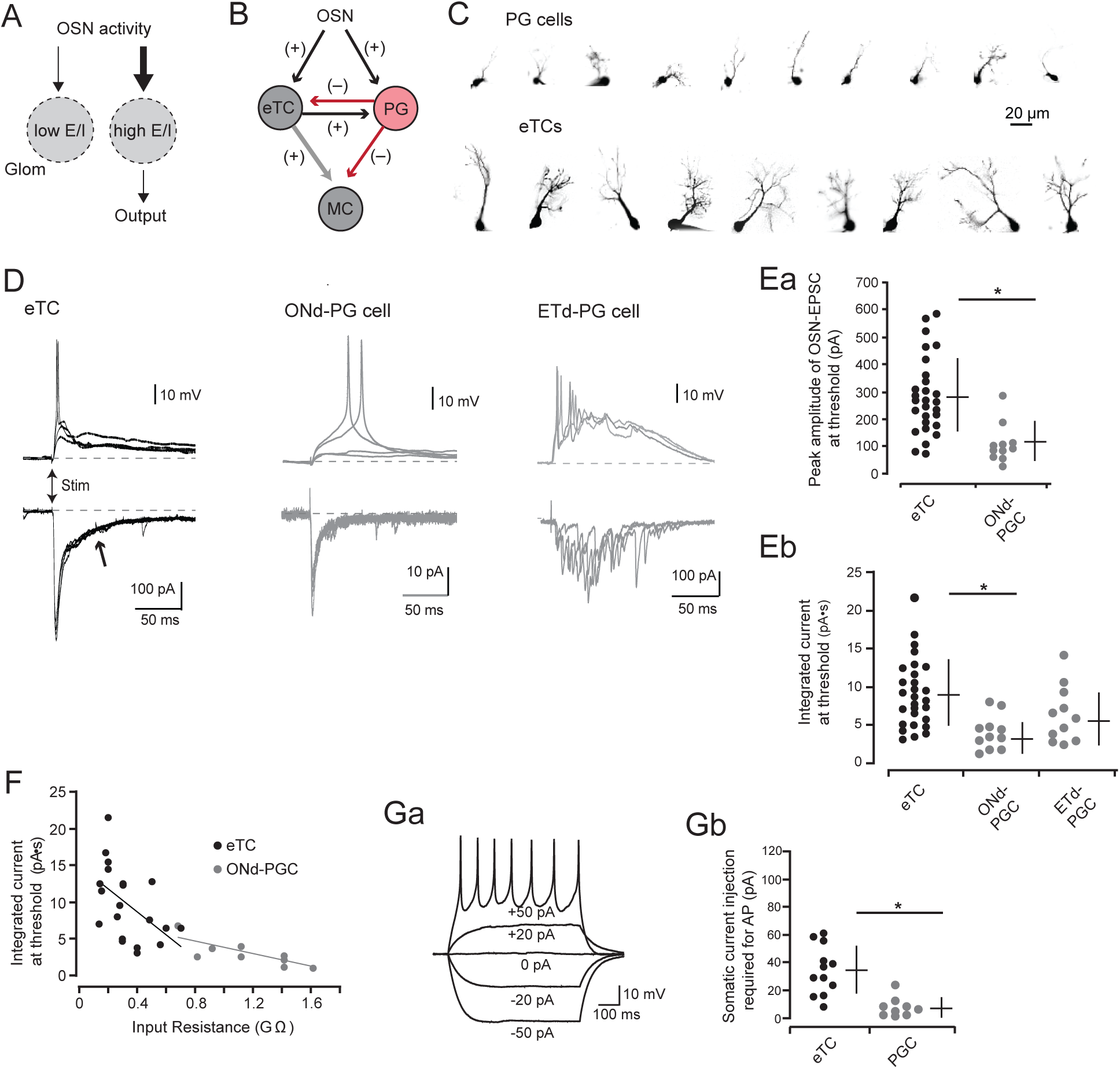
PG cells spike at lower levels of excitatory input current than eTCs. **A**, Conceptual illustration of the intraglomerular high-pass filter model. Weak olfactory sensory neuron (OSN) activity (left) results in an unfavorable excitation/inhibition (E/I) balance in a glomerulus (Glom) and block of an output, while strong OSN activity (right) produces enough excitation to overcome inhibition. **B**, Simplified circuitry of a glomerulus. OSNs can excite both glutamatergic external tufted cells (eTCs) and GABAergic periglomerular (PG) cells, which in turn excite or inhibit mitral cells (MCs). eTCs and PG cells can also reciprocally excite and inhibit each other. eTC-to-MC excitation is shown as a gray rather than black arrow to indicate a non-synaptic mechanism (Gire et al., 2012). Individual PG cells can differ in their mode of excitation (see Part **D**), which is not illustrated. **C**, Images of different Alexa-488-labeled PG cells and eTCs that were used in the electrophysiological analysis. **D**, Example current-clamp (top) and voltage-clamp (bottom; Vhold = -77 mV) recordings in response to OSN stimulation in an eTC (black traces at left), an ONd-PG cell (gray traces in middle), and an ETd-PG cell (gray traces at right). Stimulus intensities were chosen that were perithreshold for spike activity in each cell. Three or four overlaid traces for each. Note the different current response dynamics for the different cell-types, including the presence of a significant slow current in the eTC (at arrow) that occurs after the fast OSN-EPSC, as well as barrages of fast EPSCs in ETd-PG cells that arise due to polysynaptic signaling. **E**, Summary plot showing the peak amplitudes of the monosynaptic OSN-EPSCs (**Ea**) or the integrated current (**Eb**) at perithreshold stimulus intensities for eTCs and PG cells. For OSN-EPSC peak amplitude measurements, results for ETd-PGCs are not shown, since the spiking at threshold was most commonly associated with more delayed EPSCs. p < 0.0001, Mann-Whitney U test for **Ea**; p = 0.0002, Mann-Whitney U test for **Eb. F**, Plot relating the integrated current at perithreshold intensities versus the input resistance of the test ONd-PG cells or eTCs. Superimposed lines reflect linear regression fits of the eTC or PG cell data points, indicating a significant negative correlation in each case (R2 ≥ 0.27 for each; p ≤ 0.02). **G**, PG cells are more responsive than eTCs to direct current injection (600 ms) into the cell soma. Part **Ga** illustrates example voltage responses in a PG cell to currents of different amplitude, while **Gb** is a summary plot showing significant differences in the current required for spiking between eTCs and PG cells (*p = 0.0002, Mann-Whitney U-test).

While a single glomerulus operating as a high-pass filter provides an attractive, simple mechanism to contribute to stimulus contrast, there remain numerous unresolved issues. These include the fact that testing the model in any direct manner *in vivo* based on odor-evoked responses is extremely difficult. Because any one odor will activate many glomeruli, it is virtually impossible to disentangle the contribution of intraglomerular mechanisms at one glomerulus from co-occurring interglomerular mechanisms involving other bulbar components (Aungst et al., 2003; Whitesell et al., 2013; Banerjee et al., 2015) that could also contribute to stimulus contrast (Mori et al., 1999; Urban and Arevian, 2009; Tan et al., 2010; Economo et al., 2016; Zavitz et al., 2020). In addition, any understanding of the function of PG cells as a filtering mechanism must be done in the context of the newly recognized complexity of the glomerular circuitry. In terms of signaling between OSNs and output MCs, there is now excellent evidence that MCs can be excited by a feedforward pathway that involves a class of glutamatergic interneurons known as external tufted cells (eTCs, distinct from output TCs) with a dendritic arbor that is confined to one glomerulus (De Saint Jan et al., 2009; Najac et al., 2011; Gire et al., 2012). Indeed, this feedforward pathway is the dominant pathway for exciting MCs, at least as judged by the excitatory charge evoked by OSN stimulation in bulb slices (Gire et al., 2012; Vaaga and Westbrook, 2016). Because both PG cells and eTCs can both be excited by OSNs (Hayar et al., 2004a; Murphy et al., 2005) and also transmit signals to output MCs (Najac et al., 2015; Shao et al., 2019), these two cell-types arguably reside in parallel but opposing positions in the glomerular microcircuitry (**Fig. 1B**). Hence, a major part of understanding whether a glomerulus acts as a high-pass filter requires assessing the relative effectiveness of OSNs in exciting PG cells versus eTCs.

Here, we tested a number of predictions of the intraglomerular filter model using brain slice methods and weak electrical stimulation that resulted in activation of OSNs at one or at most a few glomeruli. We focused on two sets of variables in the analysis: (1) the activation probabilities for PG cells and eTCs; and (2) the inhibitory and excitatory currents that result secondarily from activation of PG cells and eTCs. The latter provided a measure of the intraglomerular E/I balance in response to fixed levels of OSN activity. We found that, in contrast to what might be predicted by their high input resistance, individual PG cells were no more likely to be excited than eTCs at low levels of OSN activity. PG cells spiked in response to much smaller synaptic currents due to their high resistance, but this advantage in terms of excitability was offset by the fact that PG cells receive far less synaptic input at equivalent levels of OSN activity. However, consistent with the predictions of a high-pass filter, we found that the intraglomerular E/I balance became markedly more favorable with increasing OSN input. We also identified factors at the level of neurotransmitter regulation that contribute to the E/I balance.

## Methods

### Ethical approval

All experiments were approved by the Institutional Animal Care and Use Committee at the University of Colorado Anschutz Medical Campus in accordance with guidelines set by the U.S. Department of Health and Human Services and outlined in the Public Health Service Policy on Humane Care and Use of Laboratory Animals. The authors understand the ethical principles under which the *Journal of Physiology* operates and our work complies with the *Journal*’s animal ethics checklist.

### Animals and slice preparation

Male and female 8-20 day-old Sprague Dawley rats obtained from Charles River Laboratories were used in most experiments. Some experiments utilized transgenic rats expressing Venus fluorescent protein under control of the vesicular γ-aminobutyric acid (GABA) transporter promoter (VGAT-Venus rats, Uematsu et al., 2008, strain 2; Wistar background; Whitesell et al., 2013). Experimental VGAT-Venus rats were heterozygotes obtained by breeding a homozygous VGAT-Venus male with a wild type Wistar female. Rats were housed with littermates (4-8 per cage) under a 12-hour light/dark cycle (lights on at 0700 hours, off at 1900 hours) with *ad libitum* access to food and water. Acute horizontal olfactory bulb slices (300-400 μm thickness) were prepared following isoflurane anesthesia and decapitation as described previously (Zak et al., 2015).

### Electrophysiological recordings

Experiments were carried out under an upright Zeiss Axioskop2 FS Plus microscope (Carl Zeiss MicroImaging) fitted with differential interference contrast (DIC) optics, video microscopy and a CCD camera (Hamamatsu). Identified cells were visualized with 10x or 40x Zeiss water-immersion objectives. Recordings were performed at 31-34°C.

The base extracellular recording solution contained the following (in mM): 125 NaCl, 25 NaHCO_3_, 1.25 NaHPO_4_, 25 glucose, 3 KCl, 1 MgCl_2_, 2 CaCl_2_ (pH 7.3 and adjusted to 295 mOsm), and was oxygenated (95% O_2_, 5% CO_2_). The pipette solution for most whole-cell recordings contained: 125 K-gluconate, 2 MgCl_2_, 0.025 CaCl_2_, 1 EGTA, 2 Na_3_ATP, 0.5 Na_3_GTP, and 10 HEPES (pH 7.3 with KOH). For the analysis of inhibitory and excitatory currents in eTCs in response to OSN stimulation, the K-gluconate in the pipette solution was replaced with an equimolar amount of cesium methanosulfonate, as well as the sodium channel blocker QX-314 (10 mM) to block action potentials. All whole-cell recordings included 100 μM Alexa 488 or Alexa 594 in the pipette solution to allow visualization of cell processes. Loose cell-attached recordings in VGAT-Venus rat slices were made with a pipette that contained the extracellular solution. Patch pipettes, fabricated from borosilicate glass, were pulled to a resistance of 4-6 MΩ for eTCs and 6-8 MΩ for PG cells. Current and voltage signals in the single- and pair-cell experiments were recorded with a Multiclamp 700B amplifier (Molecular Devices, San Jose, CA), low-pass filtered at 1.8 kHz using an eight-pole Bessel filter, and digitized at 10 kHz. Data were acquired using Axograph X software. Reported membrane potential values were corrected for a 7 mV liquid junction potential for our recordings.

Cell identity in the recordings in slices from wild-type rats was determined in part by visualizing Alexa 488- or Alexa 594-mediated fluorescence signals. eTCs were distinguished from other cells in the glomerular layer by their position in the inner half of the layer, their relatively large, spindle-shaped somata (≥10 μm diameter), a single highly branched apical dendrite that filled most of a glomerulus and no lateral dendrite, and a relatively low input resistance (∼0.2 GΩ; Hayar et al., 2004b). PG cells were identified by their small soma (≤10 μm diameter), small dendritic arbor that was confined to a part of one glomerulus, and high input resistance (> 0.5 GΩ; Puopolo and Belluzzi, 1998; McQuiston and Katz, 2001; Smith and Jahr, 2002; Hayar et al., 2004b). Fluorescence measurements were performed under whole-field epi-illumination using a DG-4 light source (Sutter Instruments, Novato, CA). Signals were detected by a CoolSnap II HQ CCD camera (Photometrics, Tucson, AZ) under control of Slidebook (Intelligent Imaging Innovations, Denver, CO) software. eTCs were further discriminated from PG cells based on their excitatory current response to OSN stimulation. eTCs, but not PG cells, displayed a large slow, monophasic current that followed the initial fast current reflecting direct OSN input. This slow current is due to recurrent excitation derived from other excitatory neurons (Gire et al., 2019). PG cells were further differentiated based on their functional responses to OSN stimulation (Shao et al, 2009): ONd-PG cells that had only a short-onset excitatory post-synaptic current reflecting direct inputs from OSNs (the OSN-EPSC) or ETd-PG cells that had a barrage of rapid EPSCs reflecting the polysynaptic OSN-to-eTC-to-PG cell pathway. In the dual-cell patch recordings, we determined that the test cells sent their apical dendrites to the same glomerulus based on anatomical measurements as well as in trial-to-trial correlations in the excitatory current responses.

Stimulation of OSN axons was performed using a broken-tip patch pipette (5-10 μm-diameter) placed in the olfactory nerve layer, 50-100 μm superficial to the glomerular layer. Current injections were delivered by a stimulus isolator (World Precision Instruments) under control of a TTL output from Axograph X software. Weak intensities of electrical stimulation were used (1 – 50 µA). This enabled us to stimulate OSNs at one glomerulus with little or no activity in neighboring glomeruli (McGann et al., 2005; Gire and Schoppa, 2009). Stimulus artifacts in most of the illustrated traces in the figures were blanked or truncated.

### Experimental Design and Statistical Methods

Data were analyzed using Axograph, Microsoft Excel, or MATLAB and are generally expressed as Mean (Standard Deviation). Significance was most commonly determined using two-tailed non-parametric tests, either the Wilcoxon matched-pairs signed-ranks test or the Mann-Whitney U test. A value of *p* < 0.05 was considered significant (single asterisks in the figures), except if multiple comparisons were made, in which case the Bonferroni correction was applied. N-values reported in the main text refer to the number of test cells, and all statistical comparisons were made based on measurements across different test cells. The numbers of rats used are also reported.

### Analysis of electrophysiological data

To evaluate the relationship between excitatory synaptic current and spiking in eTCs and PG cells, we first determined in current-clamp the OSN stimulation intensity required to cause spikes in at least half but not all trials and then measured the excitatory current in voltage-clamp. The fast sodium spikes in PG cells in particular were quite variable in amplitude (30-60 mV) and number of evoked spikes. Some PG cells did not spike at all, potentially reflecting an immature state (Benito et al., 2018); these cells were excluded from the analysis. In the analysis of the excitatory currents, we assessed both the peak amplitude of the OSN-EPSC and also the integrated current. A 300-ms window after stimulation was used for most current integrations. Excitatory currents recorded at our test holding potential (–77 mV) had negative values; however, for ease of comparison, we converted the negative currents into absolute (positive) current values.

In the analysis of the evoked excitatory synaptic currents in PG cells, synaptic delays were estimated from the time between the stimulus artifact to the start of the negative current deflection associated with the first EPSC in the response. Because of the large size of the evoked synaptic currents and low baseline noise, the start-time for the EPSCs could be determined manually with a high degree of precision. Jitter was determined from the standard deviation of the delay values from all stimulus trials at a given OSN stimulation intensity.

To estimate the single-fiber OSN-EPSC in PG cells and eTCs, we adjusted the stimulation intensity applied to OSNs to a point at which response failures were observed in most but not all trials. The mean of the OSN-EPSCs observed in successful trials at this “minimal” stimulation intensity was taken to be the single-fiber OSN-EPSC for that cell. For ETd-PG cells that had responses that included both a monosynaptic OSN-EPSC as well as polysynaptic EPSCs, a trial was considered to be a success only if it included a monosynaptic EPSC (i.e., many of the response failures for this analysis included polysynaptic EPSCs). The relative level of convergence of OSN axons onto PG cells and eTCs was estimated from values for the fiber fraction (Hooks and Chen, 2006), which was the amplitude of the single-fiber OSN-EPSC divided by the maximal OSN-EPSC evoked by strong OSN stimulation. The value for the fiber fraction for a given cell will lead to an erroneous estimate for convergence onto that cell if the release probability is less than one (as it generally is). However, a good estimate of the *relative* level of convergence onto PG cells versus eTCs can be obtained from the fiber fraction values across a population of PG cells and eTCs, since the release probabilities for OSN synapses onto PG cells and eTCs appear to be similar (Murphy et al., 2004; based on paired pulse ratios). Absolute estimates of convergence in slices were also likely to be in error due to slice artifacts. However, we assume that the fraction of OSN axons that were cut by the slicing that terminate on eTCs or PG cells was similar, enabling estimates of relative convergence onto these cells.

To estimate the E/I ratios in glomeruli, we used the approach reported in our recent paper (Gire et al., 2019) involving current recordings in eTCs. Currents at negative voltages include the OSN-EPSC and a slow recurrent excitation, while large outward currents appear at depolarized voltages that are eliminated by GABA_A_ receptor blockers that reflect inputs from glomerular layer inhibitory interneurons. The E/I ratio was determined from the ratio of the integrated recurrent excitatory conductance (*G*_*E(Recur)*_) versus the integrated inhibitory conductance (*G*_*I*_), *G*_*E(Recur)*_/ *G*_*I*_. To estimate *G*_*E(Recur)*_, we first subtracted off the OSN-EPSC from the composite excitatory current (Gire et al., 2019). In the plots the related *G*_*E(Recur)*_ and *G*_*I*_ to the amplitude of OSN-EPSC, the OSN-EPSC for *G*_*I*_ was taken from the currents recorded at negative voltages at the exact same stimulus intensity (during different trials).

### Fura-2 imaging in VGAT-Venus rat slices

Slices obtained from VGAT-Venus rats were bulk loaded with 50 μM Fura-2,AM (Invitrogen; Carlsbad, CA) 0.2% Pluronic F-127 in our standard extracellular solution but containing 2 mM MgCl_2_, 1 mM CaCl_2_ for 40 minutes in an oxygenated chamber at room temperature. Slices were then transferred into a holding chamber and allowed to recover for an additional 30 minutes at room temperature before imaging. Fluorescence signals were detected with a CoolSnap II HQ CCD camera (Photometrics; Tucson AZ) by measuring the 510 nm emission following whole-field epi-illumination at both 340 nm and 380 nm excitation wavelengths from a Lambda DG-4 light source (Sutter Instrument; Novato, CA). Data were acquired through a 40x objective as 696 × 520 pixel, 16-bit tiff images with a 20 ms exposure at 4-6 Hz using Slidebook software (3i; Denver, CO). The field of view was 225 μm × 168 μm. The images displayed in the Fura-2 measurements were further cropped to remove portions of the external plexiform layer absent of glomerular neurons.

For analysis, images were exported as raw tiff files for processing using custom MATLAB scripts (Mathworks, Natick, MA). Prior to processing, images were binned at 2×2 by averaging pixel values in each bin. For each experiment, a template image was generated from the first 200 frames in the 340 nm excitation channel. Non-rigid motion compensation was then performed (NoRMcorre; Pnevmatikakis and Giovannucci, 2017). Frame shifts were then applied to the 340 nm and 380 nm excitation channel, as well as the VGAT-Venus image. For each frame a ratiometric image was then computed by dividing emitted fluorescence signal following 340 nm and 380 nm excitation. Regions of interest (ROIs) were automatically detected from the ratiometric images using constrained nonnegative matrix factorization (CaIMaN; Pnevmatikakis et al., 2016) and were further filtered by size and shape to separate merged cells.

To analyze the responsiveness of a population of cells around a glomerulus, we determined the ratiometric Fura-2 signal (ΔR/R) within specific ROIs, then transformed them into z-scores. In the z-score calculations, at least 10 frames prior to stimulus delivery were used to estimate the mean and standard deviation of the baseline. Whether a cell was responsive was determined by averaging the z-score of three frames centered on the peak z-score found in the 20 frames post-stimulus (corresponding to a window of 5 seconds) and comparing this average value to a predetermined threshold for activity (see below). The 5-second window was long enough to account for the slow kinetics of Fura-2 response, while sufficiently short that any detected activity would be associated mainly with a primary activity event following OSN stimulation rather than secondary events that can sometimes appear many seconds after stimulation in rat OB (Schoppa and Westbrook, 2001). In the analysis, we focused on cells that lined the perimeter of the glomerulus that was closest to the stimulating electrode. We could not be certain that all of the cells were associated with the stimulated glomerulus. However, in seven cells that were whole-cell patched after the imaging to determine morphology, all seven sent apical dendrites into the glomerulus closest to the stimulating electrode.

To determine a threshold for counting a VGAT+ or VGAT-cells as responsive based on the Fura-2 signals, control experiments were performed in which visually identified VGAT+/– cells were simultaneously recorded in loose cell-attached configuration while imaging Fura-2 fluorescence responses. Stimulus intensities were adjusted to yield probabilistic single spike responses or failures. The distributions of the responses corresponding to one spike and zero spikes were clearly delineated at z-score = 2.0, and this value was used to classify responding and non-responding cells. In addition to finding criteria that separated the calcium responses based on whether there were one or zero stimulus-evoked spikes, we also calculated a distribution of the pre-stimulus noise. This was done by averaging the z-scores of three frames centered on the peak response in the 20 frames pre-stimulus.

For classifying cells as VGAT+ or VGAT–, all YFP images were normalized by dividing each pixel value by the 95^th^ quantile of all pixel intensities in a field of view. To determine whether cells contained YFP fluorescence, ROIs were constructed from neuropil regions of the image. Random pixels were selected from neuropil until the total pixel number matched the number of pixels in a given cell. The mean fluorescence signal was then computed from the pseudo-ROI. We repeated this process 10,000 times for each cell to construct a distribution of fluorescence intensities. A cell was classified as YFP-positive if its mean fluorescence exceeded 3.5 standard deviations of the bootstrapped pseudo-ROI distribution. YFP signals in cells that were counted as VGAT+ varied significantly and also displayed relative fluorescence intensities that differed significantly from the Fura-2 signals in the same cells. These variations were to be expected given differences in how these two fluorescent reporters are expressed (genetically-encoded versus acute loading). Some variations in the YFP signal in VGAT+ cells might also be explained by some cells being slightly out-of-focus. The important point however is that even these cells had YFP signals that were significantly above background, as determined by our unbiased boot-strapping approach.

### Data Availability Statement

The data that support the findings of this study are available from the corresponding author upon reasonable request.

## Results

To assess the relative effectiveness of OSNs in exciting PG cells versus eTCs, we considered that spiking in these two cells will depend not just on how easily a given synaptic current input can drive spikes (a neuron’s “excitability”) but also on *how much* synaptic input each cell receives at a given level of OSN activity. Very small neurons such as PG cells may be more excitable than larger neurons, but may receive far less synaptic input, offsetting their advantage in terms of excitability. Thus, in our comparisons between PG cells and eTCs, we measured: (1) how much synaptic current input is required to drive spiking; (2) the amount of synaptic current received by the two cell-types at equivalent levels of OSN activity; and (3) spike activity at equivalent levels of OSN activity. PG cells and eTCs were differentiated in the patch-clamp recordings based on morphological parameters such as cell size (**Fig. 1C**), as well as their excitatory current responses following OSN stimulation (**Fig. 1D**; see *Methods*). PG cells were further differentiated into functionally-defined subtypes (Shao et al., 2009): ONd-PG cells that had only a short-onset excitatory post-synaptic current reflecting direct inputs from OSNs (the OSN-EPSC) or ETd-PG cells that had a barrage of rapid EPSCs reflecting the polysynaptic OSN-to-eTC-to-PG cell pathway.

### PG cells require less synaptic current input to drive spiking than eTCs

To determine the synaptic input current level required to elicit spiking in PG cells and eTCs, we conducted a series of single-cell recordings of responses to electrical stimulation of OSNs (100 µsec; 1-50 µA). This was done by first determining in current-clamp mode the stimulation intensity that was perithreshold for spiking (≥50% spike probability; see top traces in **Fig. 1D**) and then measuring the excitatory current evoked by the same stimulus in voltage clamp (holding potential = –77 mV; bottom traces in **Fig. 1D**). We found that PG cells were much more responsive than eTCs by this measure. Comparing ONd-PG cells and eTCs, the magnitude of the OSN-EPSCs in the PG cells at perithreshold stimuli was only ∼30-40% of that in eTCs, as assessed by both peak (absolute) current amplitude (**Fig. 1Ea**; mean (SD) = 281 (135) pA for eTCs, *n* = 28 cells from 16 rats; 116 (73) pA for ONd-PG cells, *n* = 11 cells from 9 rats; *p* < 0.0001 in Mann-Whitney U test) and integrated current (**Fig. 1Eb**; 9.0 (4.3) pA•sec for eTCs, *n* = 28; 3.3 (2.2) pA•sec for ONd-PG cells, *n* = 11; *p* = 0.0002 in Mann-Whitney U test). There was also a trend for evoked synaptic currents at perithreshold stimulation intensities to be smaller in ETd-PG cells versus eTCs (integrated charge for ETd-PG cells = 5.6 (3.5) pA•sec, *n* = 11 cells from 8 rats, *p* = 0.053 in Mann-Whitney U test in comparisons with eTCs).

Our results showing that the evoked synaptic currents at perithreshold stimuli were generally smaller in PG cells than in eTCs are consistent with the much higher input resistance of PG cells (0.5-2 GΩ; Puopolo and Belluzzi, 1998; McQuiston and Katz, 2001; Smith and Jahr, 2002; Hayar et al., 2004b) versus eTCs (∼200 MΩ; Hayar et al., 2004b). These input resistance differences should naturally make PG cells more responsive to OSN input than eTCs. A mechanism based on input resistance differences was also supported by the fact that, within the population of PG cells or eTCs, the excitatory synaptic current necessary to drive spiking was negatively correlated with cell input resistance (**Fig. 1F**; ONd-PG cells: *R*^*2*^ = 0.61, *n* = 9, *p* = 0.013; eTCs: *R*^*2*^ = 0.27, *n* = 20, *p* = 0.020). In addition, we found that PG cells spiked in response to much smaller currents upon direct current injection applied to the cell soma (600-ms current steps; 34.4 (17.6) pA for eTCs, *n* = 12; 9.1 (6.5) pA for PG cells, *n* = 9; *p* = 0.0002 in Mann-Whitney U-test; **Fig. 1G**). The lower sensitivity of eTCs to OSN activation was not an artifact of the whole-cell recording condition. In a previous report (Hayar et al., 2004b), the resting potential of eTCs progressively hyperpolarized from mean values of –67 mV to –81 mV (values reported in that study were corrected here for a 9 mV junction potential) over the course of a whole-cell recording. This shift in principle could make it more difficult for these cells to reach spike threshold over time. However, the baseline potentials in our eTC recordings just prior to stimulation were relatively depolarized (mean (SD) = –67.7 (7.0) mV, *n* = 30). These results, taken together, suggest that their high input resistances enable PG cells to be excited at far lower levels of synaptic current input than eTCs.

Our recordings from ETd-PG cells revealed one other notable aspect of their physiological response to OSN stimulation. ETd-PG cells receive a polysynaptic excitatory signal derived from eTCs, but it is ambiguous whether these cells *also* receive direct monosynaptic input from OSNs. One study (Shao et al., 2009) suggested that ETd-PG cells receive “weak” OSN input, while a second study (Najac et al., 2015) suggested that there was a significant population of PG cells that displayed prolonged EPSC barrages following OSN stimulation but not monosynaptic OSN-EPSCs. In our study, we consistently observed evidence for monosynaptic OSN-EPSCs in ETd-PG cells as long as we expanded the range of OSN stimulation intensities that were investigated (**Fig. 2A-C**). In nine recordings from ETd-PG cells (across 7 rats) in which a range of stimulation intensities was sampled, weak stimuli (mean across all cells = 18 µA) typically evoked EPSCs with the relatively long onset latencies predicted by a polysynaptic mechanism (6.1 (4.5) ms), but moderate intensity stimuli (mean = 29 µA) resulted in short-onset EPSCs (1.8 (0.3) ms; *p* = 0.005 in Wilcoxon Signed-ranks test comparing EPSC delays following weak and moderate stimuli), along with a more delayed current. The early EPSCs at these moderate stimulation intensities also displayed very low jitter (mean standard deviation in onset delay = 77 (25) µsec, *n* = 9; **Fig. 2D**), similar to values observed for the OSN-EPSCs in ONd-PG cells (83 (27) µsec, *n* = 7 cells in 7 rats) at similar moderate OSN stimulation intensities (mean = 30 µA for ONd-PG cells). Importantly, the OSN-EPSCs in ETd-PG cells (264 (156) pA, *n* = 9; **Fig. 2E**) were just as large as those observed in ONd-PG cells (243 (161) pA, *n* = 7; **Fig. 2E**) at the same moderate stimulation intensities, suggesting that ETd-PG cells receive just as much OSN input as ONd-PG cells. Thus, in the rest of our analysis below, we considered ETd-PG cells to receive substantial input directly from OSNs as well as a polysynaptic pathway involving eTCs (**Fig. 2F**).

**Figure 2.**
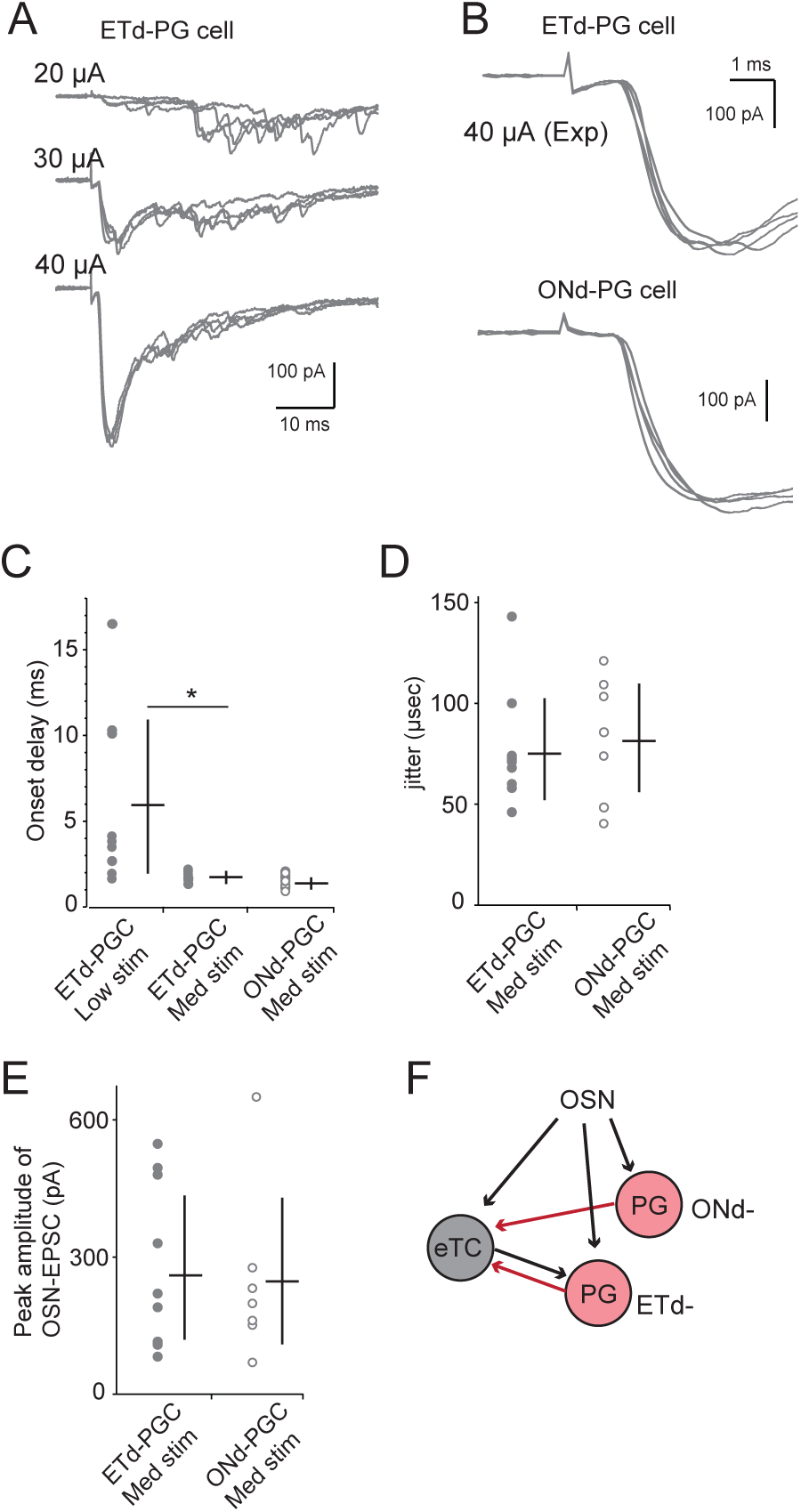
ETd-PG cells have large monosynaptic OSN-EPSCs. **A**, Current responses in an ETd-PG cell to varying intensity of electrical stimulation of OSNs (4 responses per intensity). Note that with the weakest stimuli (top) the responses were delayed, as expected for a polysynaptic OSN-to-eTC-to-PG cell mechanism, but increasing intensity stimuli recruited a fast-onset EPSC. **B**, Expanded current traces illustrating responses of an ETd-PG cell (top, 40 µA data in part A) and an ONd-PG cell (bottom; stimulus intensity = 35 µA). In each case, the EPSCs have an onset of ~1.5-2 ms following the stimulus (see artifact) and very little jitter, consistent with a monosynaptic OSN-EPSC. **C**, Summary of current onset-delay values for nine ETd-PG cell recordings and seven ONd-PG cells. For each ETd-PG cell, values are shown for weaker stimuli (mean intensity = 18 µA) or medium intensity stimuli that were ~10 µA stronger (mean = 28 µA). The stimulus intensities for ONd-PG cells (mean = 30 µA) were similar to the medium stimuli used on the ETd-PG cells. *p = 0.005, Wilcoxon matched-pairs signed rank test. **D**, Summary values for the jitter (standard deviation) in the onset delay values for each recording from ETd-PG cells and ONd-PG cells. The stimulus intensities used for this analysis were the same medium intensities shown in part **C. E**, ONd-PG cell and ETd-PG cells had similar peak amplitudes for OSN-EPSCs at medium stimulation intensities. **F**, Circuit model for ONd-PG cell and ETd-PG cells that includes the results presented here. ETd-PG cells receive substantial direct OSN input as well as a polysynaptic OSN-to-eTC-to-PG cell signal. ONd-PG cells are excited exclusively by direct OSN input.

### PG cells have much smaller excitatory currents than eTCs at a given level of OSN activity

Having determined that both ONd-PG cell and ETd-PG cells require a much lower level of excitatory synaptic current to drive spikes versus eTCs (**Fig. 1**), we next measured how much current input the different cell-types receive at a given level of OSN activity. To perform this analysis, we made simultaneous pair-cell voltage-clamp recordings in PG cells and eTCs at the same glomerulus (**Fig. 3A**). Importantly, the pair-cell configuration enabled us to compare input currents in eTCs and PG cells at the exact same level of OSN activity, not possible with single cell recordings. We confirmed that the test cells in pair-cell recordings sent their apical dendrites to the same glomerulus and thus received inputs from the same population of OSN axons based in part on anatomical measurements (see *Methods*). In addition, the PG cells and eTCs in our recordings displayed correlated excitatory current responses from trial-to-trial at the same OSN stimulation intensity (**Fig. 3B**). Such correlated responses can be explained if the two test cells were responding to activation of the same set of OSN axons that varied in their activity level from trial to trial.

**Figure 3.**
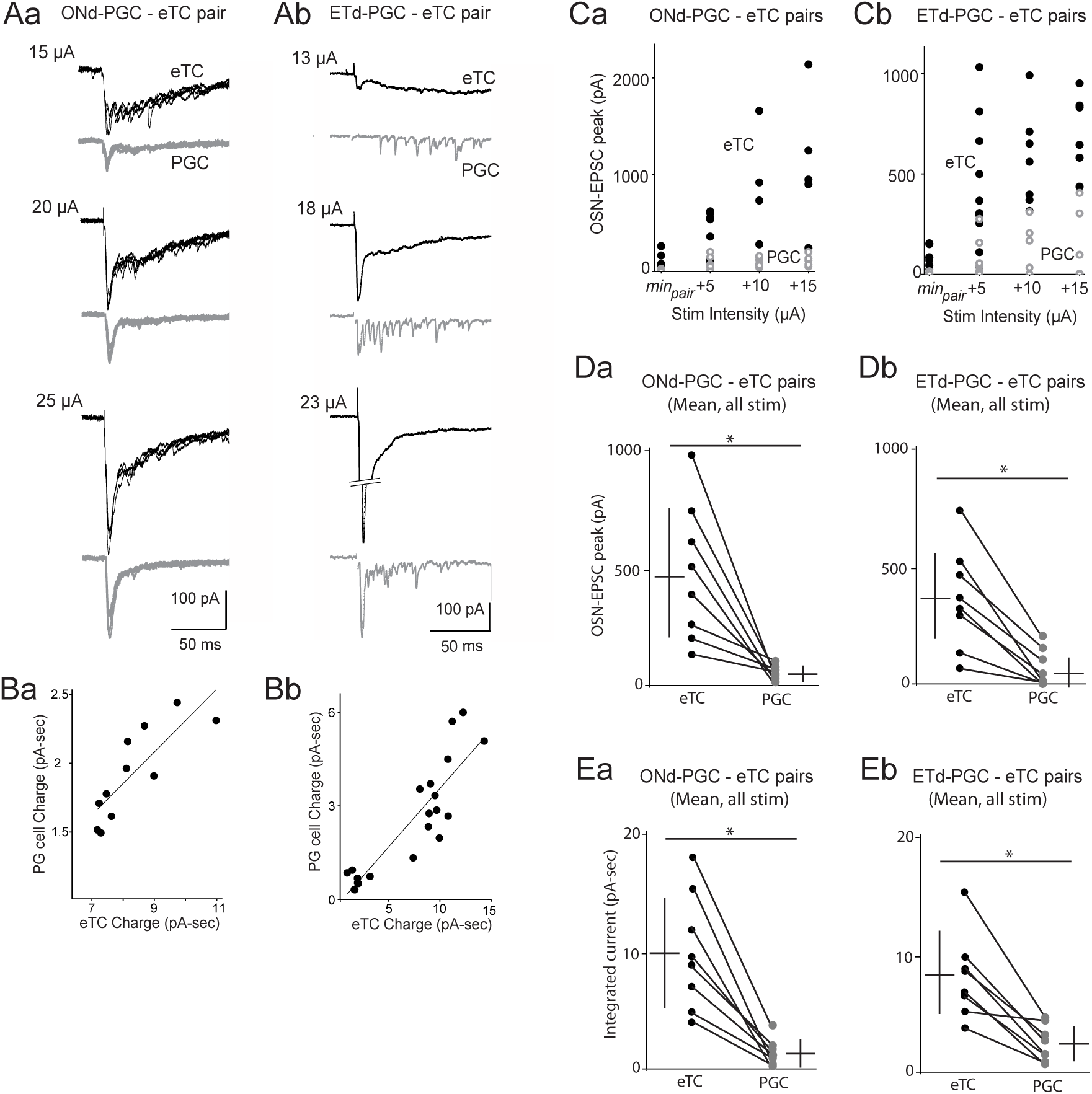
Pair-cell recordings indicate that eTCs receive far more excitatory current than PG cells at a given level of OSN activity. **A**, Example current response in pair-cell recordings that includes an eTC and either an ONd-PG cell (**Aa**) or an ETd-PG cell (**Ab**). Responses to OSN stimulation at the minimum intensity that evoked any monosynaptic OSN-EPSC in either cell (minpair) are shown at top, with responses to higher stimulation intensities shown below. **B**, Plots relating the excitatory current magnitudes in the eTC versus the PG cell across different trials at minpair for the two example pair-cell recordings in part **A** (**Ba**, 11 trials; **Bb**, 21 trials). The observed correlations across trials (p ≤ 0.002) are consistent with the two cells in each pair being affiliated with the same glomerulus (see main text). **C**, Summary of pair-cell recordings from ONd-PG cell-eTC pairs (**Ca**) and ETd-PG cell-eTC pairs (**Cb**) showing the peak amplitudes of the monosynaptic OSN-EPSCs at a range of stimulation intensities. Stimulation intensity values are shown for minpair and at indicated increments above minpair. Each data point in the two plots reflects a single recording. **D**, Summary of recordings from ONd-PG cell-eTC pairs (**Da**; n = 8) and ETd-PG cell-eTC pairs (**Db**; n = 8) in which the peak OSN-EPSC amplitude measurements across all stimulation intensities for each cell were averaged. Lines connect eTCs and PGCs from the same pairs. *p = 0.0078, Wilcoxon matched-pairs signed rank test for **Da** and **Db. E**, Summary of integrated current measurements for the same set of recordings as part **D**. As with the peak amplitudes of the OSN-EPSCs, integrated current values were consistently much larger in eTCs versus PG cells, for both ONd-PG cells (**Ea**) and ETd-PG cells (**Eb**). *p = 0.0078, Wilcoxon matched-pairs signed rank test for **Ea** and **Eb**.

In the analysis of the PG cell-eTC pairs, we considered first the amplitude of the OSN-EPSC. We found that, regardless of subtype of PG cell, the OSN-EPSCs were consistently much larger in eTCs (**Fig. 3A,C**, and **D**). Averaging across stimulus intensities, eTCs had ∼7-fold larger OSN-EPSCs in both pairs with ONd-PG cells (477 (275) pA for eTCs, 68 (32) pA for ONd-PG cells, *n* = 8 pairs in 8 rats; *p* = 0.0078, Wilcoxon matched-pairs signed rank test) and pairs with ETd-PG cells (370 (170) pA for eTCs, 55 (73) pA for ETd-PG cells, *n* = 8 pairs in 6 rats; *p* = 0.0078, Wilcoxon matched-pairs signed rank test). eTCs also had much larger excitatory currents as assessed from integrated current measurements (**Fig. 3E**). Integrated current values were ∼7-fold larger in eTCs in pairs that included ONd-PG cells (10.1 (4.5) pA-sec for eTCs, 1.4 (1.3) pA-sec for ONd-PG cells, *n* = 8; *p* = 0.0078, Wilcoxon matched-pairs signed rank test) and ∼3-fold larger in pairs that included ETd-PG cells (8.4 (3.2) pA-sec for eTCs, 2.7 (1.7) pA-sec for ETd-PG cells, *n* = 8; *p* = 0.0078, Wilcoxon matched-pairs signed rank test). Comparing ETd-PG cells and ONd-PG cells, there were more ETd-PG cells with larger integrated current values (**Fig. 3E**), as expected from the fact that these cells receive both OSN-EPSCs and polysynaptic EPSCs derived from eTCs. However, the important point was that even ETd-PG cells had much smaller excitatory currents than eTCs at a given level of OSN activity.

With the monosynaptic OSN-EPSC component of the excitatory response, it was possible to further delineate the mechanisms underlying the much larger current in eTCs. The larger OSN-EPSC could reflect larger OSN-EPSCs provided by single OSN axon fibers (single-fiber OSN-EPSCs), greater convergence of OSN axons, and/or differences in the glutamate release probability at OSN synapses. Because there is evidence that OSN synapses onto tufted cells and PG cells have similar release probabilities (Murphy et al., 2004; based on paired pulse ratios), we focused on the first two mechanisms. To determine whether the single-fiber OSN-EPSCs are different in eTCs versus PG cells, we altered our analysis of each pair-cell recording by identifying minimal stimulus conditions for both the PG cell and the eTC, i.e., stimuli at which failures were evoked in most but not all trials (**Fig. 4Aa, Ab**). Under this condition, the evoked OSN-EPSC can be assumed to reflect the input derived from a single activated OSN axon. Because of the added level of difficulty of this analysis – requiring reliable measurements of single-fiber OSN-EPSCs in both the eTC and PG cell in each pair – our yield was somewhat lower, and so we combined the results from 10 pair-cell recordings (across 8 rats) in which the test PG cell was either ONd- or ETd-. This showed that the single-fiber OSN-EPSCs were ∼2-fold larger in the eTC versus the PG cell (**Fig. 4B**; 114 (51) pA in the eTC, 50 (21) pA in the PG cell, *n* = 10; *p* = 0.0488, Wilcoxon matched-pairs signed rank test).

**Figure 4.**
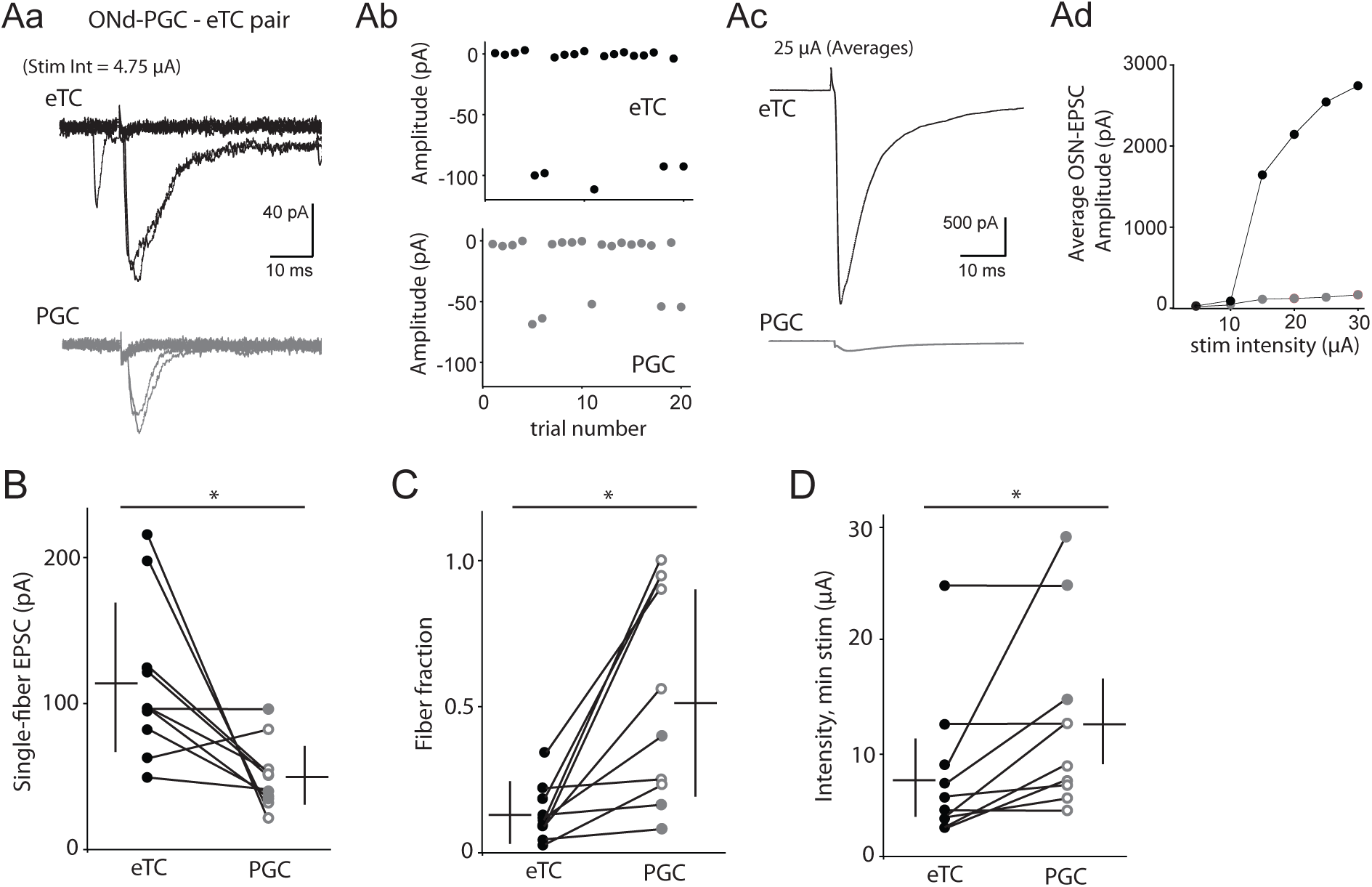
eTCs have both larger single-fiber OSN-EPSCs and greater convergence of OSN axons versus PG cells. **A**, Example recording from an ONd-PG cell-eTC pair indicating method of analysis. **Aa** and **Ab**, We first determined the amplitude of the single-fiber OSN-EPSC for each cell (eTC = –98 pA; PGC = –54 pA) from currents evoked at minimal intensity stimuli (4.75 µA) that resulted in response failures in most trials. Example current responses (**Aa**, 8 trials) and OSN-EPSC amplitudes from 20 trials (**Ab**) are shown. **Ac** and **Ad**, Relative convergence of OSN axons onto the eTC versus the PG cell was then estimated from OSN-EPSCs evoked by much stronger stimulation (25 µA). The amplitude of the single-fiber OSN-EPSC divided by this OSN-EPSC yielded a value for the fiber fraction (eTC = 0.043; PGC = 0.25), the inverse of which across a population of cells provided an estimate of relative convergence (Hooks and Chen, 2006). Average currents at 25 µA (**Ac**) and amplitudes across different stimulation intensities (**Ad**) are illustrated. **B**, Across 10 eTC-PG cell pair recordings, the single-fiber OSN-EPSC was ~2-fold larger in the eTC versus the PG cell. *p = 0.0488, Wilcoxon matched-pairs signed rank test. Results from recordings from ONd-PG cells (open symbols) and ONd-PG cells (closed symbols) were pooled. **C**, Values for the fiber fractions from the same eTC-PG cell pairs in part **B**. Note that the fiber fractions were consistently much smaller for eTCs, implying greater convergence of OSN axons. *p = 0.0020, Wilcoxon matched-pairs signed rank test. **D**, In the same 10 pairs, the minimal stimulation intensity required to elicit an OSN-EPSC was typically lower for the eTC versus the PG cell (*p = 0.0156, Wilcoxon matched-pairs signed rank test), also consistent with a greater number of OSN axons terminating on the eTC.

To test whether eTCs and PG cells also differ in their level of convergence from OSN axons, we analyzed OSN-EPSCs evoked by strong stimuli in the same 10 pair-cell recordings (example in **Fig. 4Ac,Ad**). Dividing the amplitude of the single-fiber OSN-EPSC by the OSN-EPSC evoked by strong stimuli yielded a value for the fiber-fraction for that cell (Hooks and Chen, 2006). The inverse of the mean values of the fiber fractions across a population of eTCs and PG cells provided an estimate for the relative level of convergence onto the two cell-types (since release probabilities are similar; see above). Fiber fraction values were found to be 3-to-4-fold larger in PG cells versus eTCs (**Fig. 4C**; mean fiber fraction = 0.148 (0.093) for eTCs and 0.544 (0.348) for PG cells, *n* = 10 pairs across 8 rats; *p* = 0.0020 in Wilcoxon matched-pairs signed rank test), implying that there are 3-to-4 times as many OSN axons targeting eTCs versus PG cells. That there is much more convergence of OSN axons onto eTCs versus PG cells was also supported by the values for the minimal stimulus intensities that were used to elicit the single-fiber OSN-EPSCs. Across the 10 eTC-PG cell pairs, there was sometimes no difference in the minimal stimulus intensities between the eTC and PG cell (*n* = 3 pairs, including example in **Fig. 4A**), but, when there was a difference, the eTC always displayed the lower minimal stimulus intensity (**Fig. 4D**; mean difference across 10 pairs = 5.2 (5.9) µA; *p* = 0.0156 in Wilcoxon matched-pairs signed rank test). A lower value for the minimal stimulus intensity implied that OSN axons terminating on eTCs were at a higher density than axons terminating on PG cells at or near the stimulation electrode. These results taken together indicate that eTCs have much larger OSN-EPSCs than PG cells at a given level of OSN activity, and that the larger OSN-EPSCs in eTCs likely reflect both a larger OSN-EPSC evoked by single OSN axons and also greater convergence.

### Individual eTCs and PG cells have similar spike probabilities at low OSN activity levels, but more PG cells spike due to their higher number

Our results thus far have indicated that PG cells are more excitable than eTCs to a given level of current input, but that, for a given level of OSN activity, eTCs receive much more excitatory current input. We next assessed what these competing properties meant for the spike responses of the two cell-types for given level of OSN activity. Do PG cells preferentially spike at low levels of OSN activity due to their high input resistance or does the greater input current in eTCs make them more responsive?

To address this question, we first recorded spike activity in pairs of eTCs and PG cells at the same glomerulus (**Fig. 5A**), comparing the values for the perithreshold stimulus intensities for spiking (*PT*_*spike*_, in µA) in the two cells. The cell with the lower value for *PT*_*spike*_ could be assumed to be the cell that required less OSN activity to drive spiking. Across 12 pair-cell recordings in 10 rats, we in fact found no significant differences in *PT*_*spike*_ between eTCs and PG cells (pooled ONd- and ETd-; **Fig. 5B**; 10.6 (4.4) µA for eTCs, 14.8 (4.6) µA for PG cells; *p* = 0.078 in Wilcoxon matched-pairs signed rank test). Differences in *PT*_*spike*_ were observed in seven of the pairs, but the value was smaller in the eTC in six of these. This implied that any potential trend in *PT*_*spike*_ differences between eTCs and PG cells was the opposite of that predicted by prior studies that suggested that PG cells should be preferentially excited by weak inputs due to their high input resistance (Cleland and Sethupathy, 2006; Gire and Schoppa, 2009; Fukunaga et al., 2014). The shapes of the entire stimulus-spike response curves for eTCs and PG cells were also generally quite similar (**Fig. 5C**), with both cells displaying steep curves that typically saturated at spike probabilities of 1.0. That PG cells and eTCs spiked in response to similar levels of OSN activity was also supported by the large number of single cell recordings in eTCs and PG cells from earlier in the study that related the OSN-EPSCs to spiking (**Fig. 1D**). In these experiments, the values for *PT*_*spike*_ were indistinguishable between eTCs, ONd-PG cells, and ETd-PG cells (**Fig. 5D**; 21 (12) µA for eTCs, *n* = 28; 23 (10) µA for ONd-PG cells, *n* = 11; 18 (10) µA for ETd-PG cells, *n* = 13; *p* = 0.52 in Kruskal-Wallis test). In addition, we found no relationship between *PT*_*spike*_ and the input resistance of the PG cell in the single-cell recordings (**Fig. 5E;** *R*^*2*^ = 0.06, *p* = 0.76). This argued against the possibility that there was a population of very high input resistance PG cells that were recruited at the lowest level of OSN activity.

**Figure 5.**
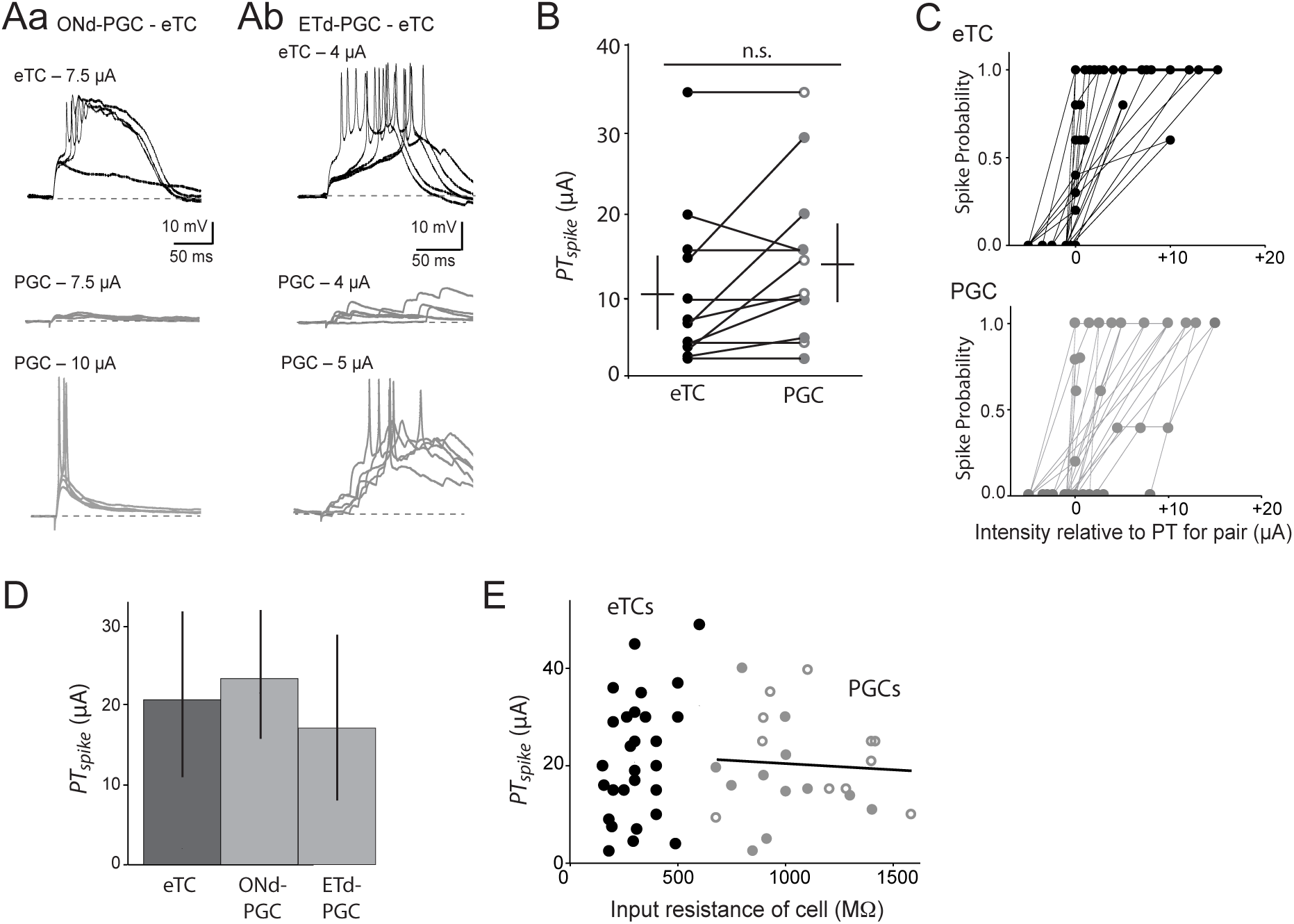
Pair-cell recordings indicate that weak OSN stimulation activated individual PG cells and eTCs at similar probabilities. **A**, Example voltage responses in pair-cell recordings that included an eTC and either an ONd-PG cell (**Aa**) or an ETd-PG cell **Ab**). In both pairs, the eTC (top traces) spiked at slightly lower stimulation intensities. For the PG cells, traces are shown both at the stimulation intensity that effectively elicited spikes in the eTC (middle traces) and at a higher stimulation intensity that elicited spikes in the PG cell (bottom traces). **B**, Summary of analysis of perithreshold stimulation intensities for spiking (PTspike) of eTCs and PG cells from 12 pair-cell recordings. In the seven pairs in which differences in PTspike values were observed between the cells, the eTC spiked at a lower intensity in six of them, although the difference was not significant across the population of recordings (p = 0.078, Wilcoxon matched-pairs signed rank test). **C**, Summary from the same 12 pair-cell recordings plotting spike probability versus stimulation intensity for eTCs (top) and PG cells (bottom). Stimulation intensity values are shown relative to that which was perithreshold for evoking spikes in at least one of the two cells in the pair. **D**, Values for PTspike derived from single-cell recordings from eTCs, ONd-PG cells, and ETd-PG cells were not significantly different from each other (p = 0.52, Kruskal-Wallis test). Similar experiments as Fig. 1Eb. **E**, Values for PTspike from the single-cell recordings for eTCs, ONd-PG cells, and ETd-PG cells plotted as a function of the input resistance of the test cell. Values for PG cells (both ONd- and ETd-) were fitted with a line, showing that there was no correlation between input resistance and PTspike (R2 = 0.06, p = 0.76).

As a second approach to evaluate the spike probability of eTCs and PG cells, we performed Fura-2AM-based calcium imaging. This technique enabled us to examine the responsiveness of a population of cells at a glomerulus to OSN stimulation at once. To help differentiate PG cells and eTCs in the imaging, we made slices from vesicular GABA transporter (VGAT)-Venus transgenic rats (Uematsu et al., 2008; **Fig. 6A**), in which GABAergic cells in the glomerular layer of the bulb are labeled in a highly specific fashion (Whitesell et al., 2013). Fura-2 appeared to be a reliable indicator of cellular activation, at least at the level of single action potentials for both VGAT+ and VGAT– cells. When we recorded both Fura-2-mediated fluorescence transients and electrical activity in response to OSN stimulation at the same time (in loose cell-attached, LCA, configuration; **Figure 6B,C**), fluorescence signals associated with single spikes were consistently much larger than when there were no spikes. Computing a Z-statistic from stimulus-evoked responses in five VGAT+ and four VGAT– cells, we found that a z-score = 2.0 perfectly delineated calcium responses that were associated with zero versus single spikes. A z-score = 2.0 was then used as the threshold to delineate responsive versus unresponsive VGAT+ and VGAT– cells in the subsequent analysis of cell counts at different OSN stimulation intensities.

**Figure 6.**
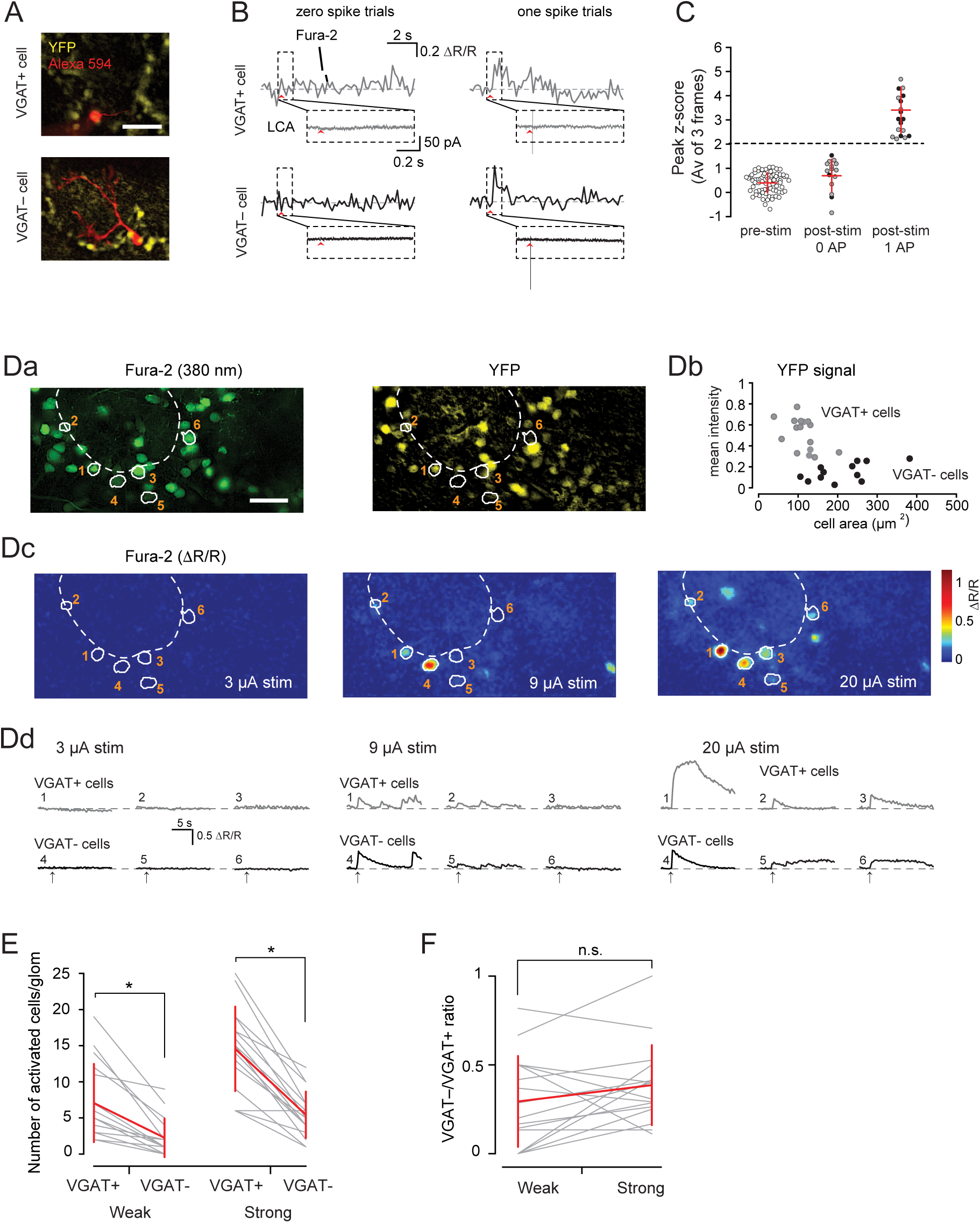
Counts of activated PG cells and eTCs using calcium imaging in VGAT-Venus rats. **A**, Example images of olfactory bulb slices taken from VGAT-Venus rats. Shown are a population of VGAT-Venus+ cells (yellow) surrounding a glomerulus along with one VGAT+ cell (top) or one VGAT– cell (bottom) that were patched and filled with Alexa-594. Scale bar = 50 µm. **B**, Example simultaneous measurements of Fura-2-mediated 340/380 ratio (DR/R) changes and electrical responses that were used to determine the threshold for counting cells as activated in the subsequent analysis in parts **D**-**F**. Responses to OSN stimulation (at arrows) are shown for one VGAT-Venus+ cell (top) and one VGAT-Venus– cell (bottom), for trials in which OSN stimulation evoked either zero (left) or single spikes (right). Spikes were recorded in the loose cell-attached (LCA) patch configuration. **C**, Summary of dual calcium-imaging and LCA patch recordings from five VGAT+ cells (gray symbols) and four VGAT– cells (black symbols) showing that the calcium responses associated with single spikes were always greater than the no-spike condition. Data reflect pooled calcium response measurements, transformed into z-scores (see Methods), with each point corresponding to one trial. Dashed horizontal line reflects the threshold (z-score = 2) used in the analysis of cell counts. **D**, Example experiment in which the number of responsive VGAT-Venus+ and VGAT-Venus– cells were counted at one stimulated glomerulus (indicated by dashed curve; stimulation electrode was above field of view). Illustrated are Fura-2 signals (380 nm excitation; **Da**, left); YFP fluorescence (**Da**, right); YFP fluorescence measurements from 30 cells associated with the glomerulus that were determined to be VGAT+ or VGAT– (**Db**); heat maps of Fura-2 fluorescence ratio changes following OSN stimulation at three stimulus intensities (3, 9, and 20 µA; **Dc**); and the time-courses of fluorescence changes in three VGAT- and three VGAT+ cells (**Dd**). Based on the Fura-2 measurements in **Dd** (5-sec window after stimulation), none of the six cells were determined to respond to 3 µA stimulation, four cells (cells 1,2,4, and 5) responded to 9 µA stimulation, and all cells responded to 20 µA stimulation. In **Db**, the y-axis reflects pixel values that were normalized by the 95th quantile of all pixel intensities in a field of view. **E**, Summary of imaging experiments conducted across 16 glomeruli in 11 rats. Plotted are the number of VGAT+ and VGAT– cells that were activated under two stimulus conditions, either weak intensity stimulation that was just above threshold for eliciting responses in the cells (left; 9 µA in example in part **D**) or strong stimulation (right; 20 µA in example in part **D**). *p < 0.0001, Wilcoxon matched-pairs signed rank test. **F**, Ratio of activated VGAT– versus VGAT+ cells under weak versus strong stimulus conditions for the same 16 glomeruli as in **E**. The ratios were not significantly different.

To test whether VGAT+ or VGAT– cells were preferentially excited at low levels of OSN activity with the imaging, we adjusted the OSN stimulation intensity until we found a level of neural activity at a glomerulus that was just reliably detected and then counted the number of activated VGAT+ and VGAT– cells based on the calculated z-scores (data at 9 µA in the example in **Fig. 6D**). At this OSN activity level, we found that there were ∼3 times more activated VGAT+ cells than VGAT– cells (**Fig. 6E;** 7.1 (5.4) VGAT+ cells and 2.3 (3.4) VGAT– cells per glomerulus; *n* = 16 glomeruli in 11 rats; *p* < 0.0001 in Wilcoxon matched-pairs signed rank test).

The ∼3-fold higher number of activated VGAT+ versus VGAT– cells with weak stimuli in the imaging experiments might appear to be inconsistent with the electrophysiological analysis that suggested that individual eTCs and PG cells are activated with similar probabilities with weak stimuli (**Fig. 5**). However, an additional variable affecting the cell counts in the imaging is that the total numbers of VGAT+ versus VGAT– cells likely are not the same. Indeed, estimates of the number of GABAergic neurons in the glomerular layer are quite high, near ∼400,000 (Parrish-Aungst et al., 2007), which compares to a value near 100,000 for the estimated number of tufted cells (Shepherd et al., 2004), which are inclusive of eTCs. To further evaluate the possibility that the greater number of VGAT+ cells with weak stimuli reflected a greater cell number in our experiments, we analyzed calcium responses in VGAT+ and VGAT– cells under conditions in which OSNs were stimulated at much higher intensities (20 µA in the example in **Fig. 6D**). We took the cell counts with these stimuli as an estimate of the number of VGAT+ and VGAT– cells at a glomerulus that could be activated and detected with our electrical stimulation and imaging methods. As expected, stronger stimuli elicited many more activated VGAT+ and VGAT-cells than weak stimuli (**Fig. 6Dd, E**), but the ratio of activated VGAT+ versus VGAT– cells was unchanged (**Fig. 6F**; VGAT–/VGAT+ ratio = 0.29 (0.24) for strong stimuli versus 0.39 (0.24) for weak stimuli, *n* = 16; *p* = 0.11 in Wilcoxon matched-pairs signed rank test). The similar activation ratios under the two stimulus conditions are consistent with the greater number of activated VGAT+ cells with weak stimuli reflecting a higher cell number rather than differences in activation probabilities.

### Impact of glutamate transporters and metabotropic glutamate receptors on the intraglomerular E/I balance

In the final section of this study, we returned to the intraglomerular high-pass filtering model proposed at the start of the study (**Fig. 1A**) in which the output of a glomerulus is selectively blocked at low OSN activity levels due to an unfavorable glomerular E/I balance. Our analysis thus far indicating that individual PG cells and eTCs have similar spike probabilities in response to synaptic input argues against at least one potential mechanism of producing an unfavorable E/I balance with low OSN activity, involving the high input resistance of PG cells (Cleland and Sethupathy, 2007; Gire and Schoppa, 2009; Fukunaga et al., 2014). Might there be other mechanisms that can contribute to condition-dependent changes in the E/I balance suitable for an intraglomerular high-pass filter?

To address this question, we turned to an approach recently reported (Gire et al., 2019) involving voltage-clamp recordings from eTCs at different voltages (**Fig. 7A**). The eTC current response to OSN stimulation at negative voltages (*V*_*hold*_ = –77 mV) includes both a fast OSN-EPSC along with a slow recurrent excitatory component (*I*_*E(Recur)*_) that arises secondarily due to spike activity in other eTCs (and possibly TCs or MCs), while recordings at depolarized voltages (*V*_*hold*_ = +28 mV) reveal a GABA_A_ receptor-mediated inhibitory current (*I*_*I*_; Shao et al., 2012) that is likely to be mainly derived from PG cells. Based on these current measurements, we assessed the integrated conductances *G*_*E(Recur)*_ and *G*_*I*_ along with the conductance ratio *G*_*E(Recur)*_/*G*_*I*_. Then, by relating *G*_*E(Recur)*_/*G*_*I*_ to the OSN-EPSCs measured at different stimulation intensities in the same recording, we estimated how the intraglomerular E/I ratio changes across different relative levels of OSN activity.

**Figure 7.**
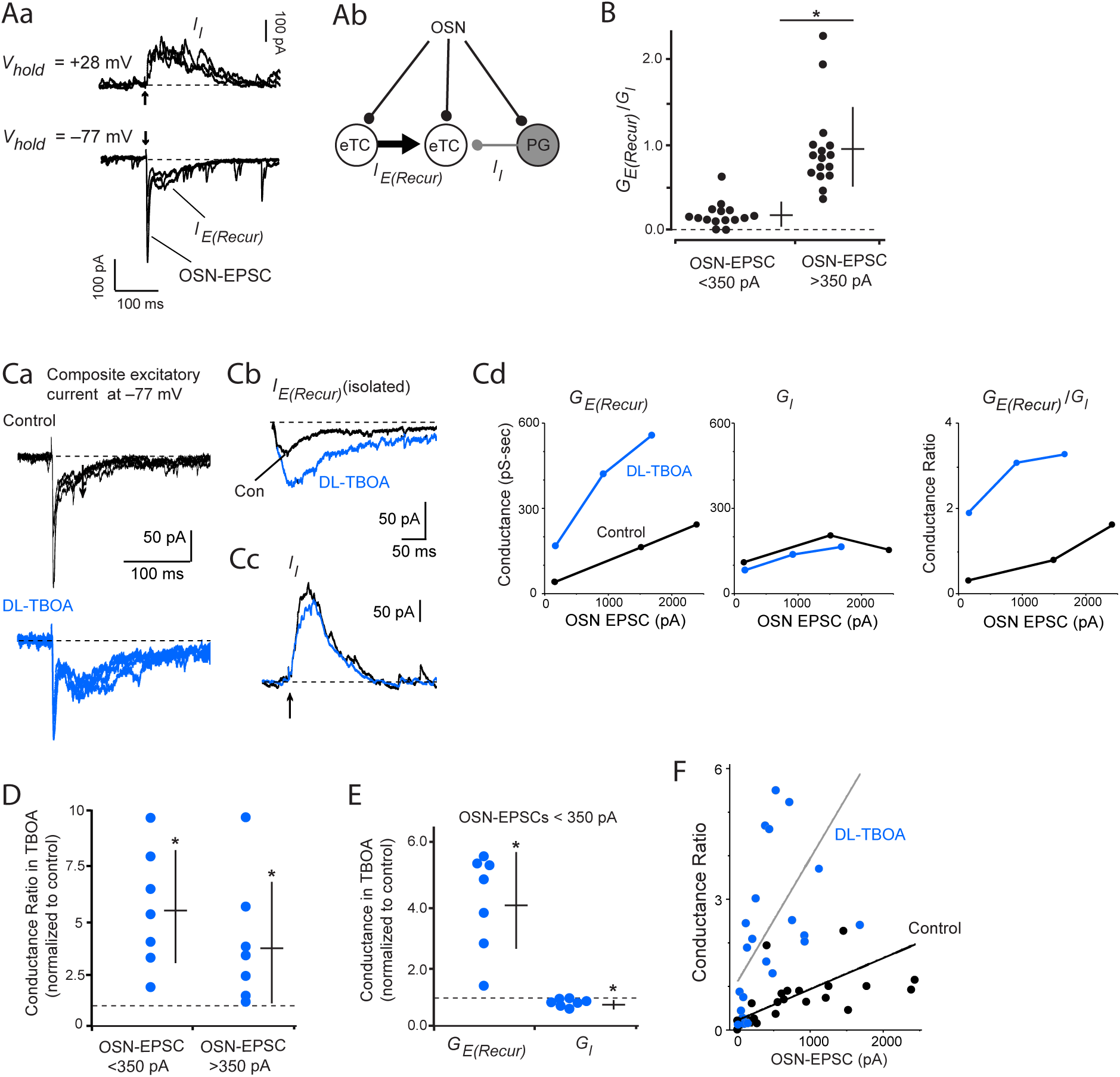
Excitation/Inhibition ratios in glomeruli as a function of OSN input: Contribution of glutamate transporters. **A**, Method of deriving E/I ratios as a function of OSN input levels. An example recording of currents evoked by OSN stimulation at different voltages in an eTC (**Aa**) and a simplified circuit representing the underlying mechanisms (**Ab**) are illustrated. At negative voltages, eTC currents include the fast OSN-EPSC along with slower recurrent excitation IE(Recur) (Gire et al., 2019). At positive voltages, an inhibitory outward current (II) appears that is likely mainly due to inputs from PG cells. **B**, Summary. At weak OSN input levels (OSN-EPSCs < 350 pA), the ratio of the integrated recurrent excitatory conductance (G_E(Recur)_) versus integrated inhibitory conductance (GI) was small (mean = ~0.20), but became much larger (mean = ~1) at higher levels of OSN activity (OSN-EPSCs > 350 pA). *p < 0.0001, Mann-Whitney U-test comparing ratios for small versus large OSN-EPSCs. Each data point reflects a single eTC recording. **C**, Effect of the glial glutamate uptake blocker DL-TBOA (10 µA) on recurrent excitation and inhibition in an example eTC recording. Illustrated are the two-component excitatory currents (**Ca**; four trials overlaid), I_E(Recur)_ isolated using a subtraction protocol (**Cb**), I_I_ (**Cc**), and the derived values for G_E(Recurr)_, G_I_, and the integrated conductance ratio (G_E(Recur)_/G_I_) plotted as a function of the OSN-EPSC amplitude for this experiment (**Cd**). **D**, Across seven eTC recordings, DL-TBOA consistently increased G_E(Recur)_/G_I_ at both weak and strong OSN input levels. Values plotted are the conductance ratios measured in DL-TBOA normalized to control ratios. *p = 0.0156, Wilcoxon matched-pairs signed rank test for both. Horizontal dashed line demarcates 1.0, i.e., no change. **E**, In the same seven eTC recordings, DL-TBOA caused a large increase in G_E(Recur)_ and a very small (but significant) decrease in G_I_. This indicated that most of the change in the conductance ratio in Part **D** was due to drug effects on G_E(Recurr)_. *p = 0.0156, Wilcoxon matched-pairs signed rank test for both. **F**. Linear regression fits of pooled data obtained from seven eTC recordings relating the conductance ratios G_E(Recur)_/G_I_ to OSN-EPSC amplitude obtained at different OSN stimulation intensities. DL-TBOA increased the slope of the relationship (p < 0.001 from ANCOVA), consistent with DL-TBOA increasing the conductance ratios irrespective of OSN input level.

As previously reported (Gire et al., 2019), *G*_*E(Recur)*_/*G*_*I*_ varied dramatically as a function of the OSN-EPSC. *G*_*E(Recur)*_/*G*_*I*_ had very small values for weak OSN-EPSCs (*G*_*E(Recur)*_/*G*_*I*_ = 0.20 (0.15) for OSN-EPSCs < 350 pA, *n* = 15 eTC recordings in 11 rats; **Fig. 7B**) but approached unity for large OSN-EPSCs (*G*_*E(Recur)*_/*G*_*I*_ = 0.96 (0.48) for OSN-EPSCs > 350 pA, *n* = 16 eTC recordings in 11 rats; *p* < 0.0001 in Mann-Whitney U test). Importantly, the observed changes in *G*_*E(Recur)*_/*G*_*I*_ as a function of the OSN-EPSC fit with the intraglomerular filter model that predicts that increasing OSN input should increasingly favor excitation (**Fig. 1A**).

We reasoned that one potential mechanism that contributes to an unfavorable E/I balance at weak levels of OSN activity is the activity of glutamate transporters. Because recurrent excitation within a glomerulus is mediated by extrasynaptic glutamate receptors (Gire et al., 2019), it should be especially susceptible to the activity of glutamate transporters that restrict the spread of glutamate in the extrasynaptic space. Indeed, consistent with this hypothesis, we found that blocking glial glutamate transporters with DL-threo-beta-benzyloxyaspartate (DL-TBOA, 10 µM) greatly increased *G*_*E(Recur)*_/*G*_*I*_ for smaller OSN-EPSCs (459 (256)% increase for OSN-EPSCs < 350 pA; *n* = 7 eTC recordings in two rats; *p* = 0.0156 in Wilcoxon matched-pairs signed rank test; **Fig. 7C-D**). This increase in the conductance ratio was mainly due to the effect of DL-TBOA on *G*_*E(Recur)*_ (310 (147)% increase, *n* = 7; *p* = 0.0156 in Wilcoxon matched-pairs signed rank test; **Fig. 7E**); the drug had only a small effect on *G*_*I*_ (20 (14)% decrease, *n* = 7; *p* = 0.0156). The effects of DL-TBOA were however not limited to conditions of low OSN input levels, as DL-TBOA also increased *G*_*E(Recur)*_/*G*_*I*_ when OSN-EPSCs were large (**Fig. 7Cd, D**; 293 (287)% increase for OSN-EPSCs > 350 pA, *n* = 7; *p* = 0.0156 in Wilcoxon matched-pairs signed rank test). That the DL-TBOA-induced increase in conductance ratios was not specific to conditions of weak input was also supported by the fact that the drug increased the slope of the lines fitted to plots that related the conductance ratios to OSN-EPSC amplitude (**Fig. 7F**). Thus, while glutamate transporter activity contributed to an unfavorable E/I balance at weak OSN input levels, saturation of these same transporters did not account for the more favorable E/I balance at high OSN input conditions.

As one mechanism that could account for the more favorable *G*_*E(Recur)*_/*G*_*I*_ conductance ratios at high OSN input levels, we considered the role of Group II metabotropic glutamate receptors (mGluRs) that are on PG cells (Ohishi et al., 1993; Sahara et al., 2001). Because Group II mGluRs can mediate a reduction in GABA release from PG cells (Zak et al., 2015), these mGluRs could at least partially account for the higher conductance ratios under strong OSN input conditions.

Consistent with this hypothesis, we found that the Group II mGluR antagonist LY341495 (1 µM) decreased *G*_*E(Recur)*_/*G*_*I*_ for large OSN-EPSCs (53 (27)% decrease in *G*_*E(Recur)*_/*G*_*I*_ for OSN-EPSCs > 350 pA, *n* = 7; *p* = 0.0156, Wilcoxon matched-pairs signed rank test; **Fig. 8A,B**). The effect of LY341495 at these OSN input levels could be attributed entirely to its effect on PG cell-mediated inhibition, since the drug consistently increased *G*_*I*_ (165 (174)% increase, *n* = 7, *p* = 0.0156, Wilcoxon matched-pairs signed rank test; **Fig. 8C**) without impacting *G*_*E(Recur)*_ (4 (20)% decrease, *n* = 7). Importantly, we found that LY341495 did not impact the conductance ratio when OSN input levels were much lower (19 (44)% decrease in *G*_*E(Recur)*_/*G*_*I*_ for OSN-EPSCs < 350 pA, *n* = 8; *p* = 0.383 in Wilcoxon matched-pairs signed rank test; **Fig. 8B**).

**Figure 8.**
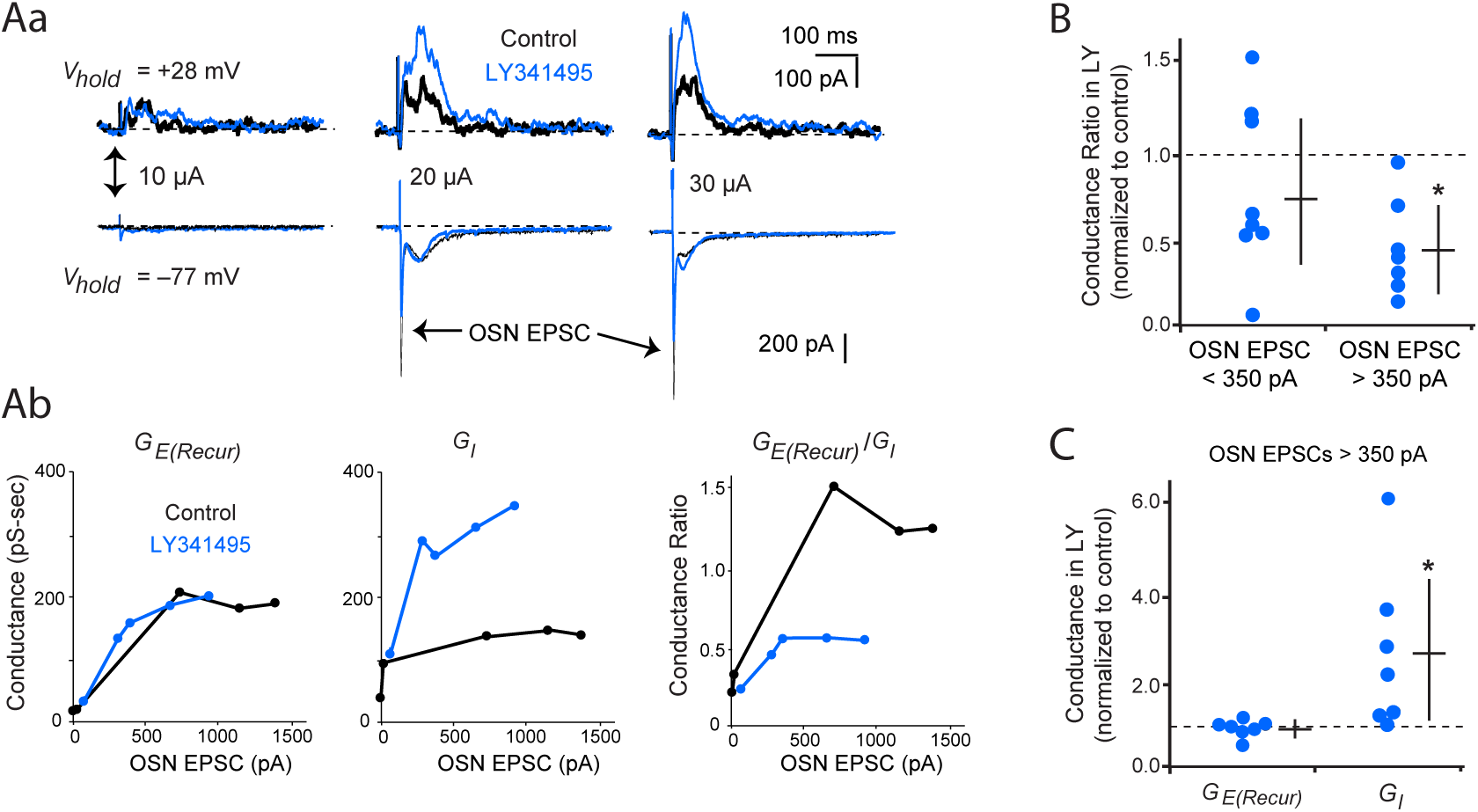
Group II mGluRs contribute to more favorable E/I ratios at high OSN input levels. **A**, Example recording of excitatory and inhibitory currents in an eTC (see Fig. 7A) testing the effect of the Group II metabotropic glutamate receptor (mGluR) antagonist LY341495 (1 µM). Illustrated are data traces (**Aa**) as well as the derived values for G_E(Recur)_, G_I_, and the conductance ratio (G_E(Recur)_/G_I_) plotted as a function of the OSN-EPSC amplitude for this experiment (**Ab**). For the data traces, averages of five trials rather than individual trials are shown to better illustrate the effects of LY341495. Note that for 10 µA OSN stimulation, there was no OSN-EPSC in the test eTC, although there was a small inhibitory current, likely derived from PG cells that received OSN input at this weak intensity. **B**, Across multiple eTC recordings, LY341495 decreased G_E(Recur)_/G_I_ at strong OSN input levels (OSN-EPSC > 350 pA; n = 7) but not weak input levels (OSN-EPSC < 350 pA; n = 8). *p = 0.0156, Wilcoxon matched-pairs signed rank test. **C**, At high OSN input levels (OSN-EPSC > 350 pA), DL-TBOA caused a large increase in G_I_ without affecting G_E(Recur)_. This indicated that the change in the conductance ratio at high OSN input levels was due to drug effects on G_I_. *p = 0.0156, Wilcoxon matched-pairs signed rank test.

Taken together, our pharmacological analyses identified two mechanisms that contribute to shaping the E/I ratios at different levels of OSN activity: glial glutamate transporters that contribute to low E/I ratios at all OSN input levels and Group II mGluRs that specifically promote high E/I ratios at high OSN input levels. These results will be discussed below in the broader context of various potential mechanisms that could shape the intraglomerular E/I balance.

## Discussion

PG cells in the olfactory bulb are proposed to mediate an intraglomerular high-pass filter through the GABAergic synapses that they target onto output MCs and TCs (Cleland and Sethupathy, 2006; Gire and Schoppa, 2009; Fukunaga et al., 2014). With this filtering mechanism, weak stimulation of OSNs (e.g., due to an odor with a low-affinity for an OR) mainly produces PG cell-driven inhibition, blocking an output, but strong stimulation generates enough excitation within a glomerulus to overcome this inhibition (**Fig 1A**). In this study, we used electrophysiological and imaging methods in rat olfactory bulb slices to test a number of predictions of this high-pass filtering model, including the hypothesis that PG cells are preferentially excited by weak stimuli due to their high input resistance. This privileged position for PG cells in terms of activation could be important for having weak stimuli produce an unfavorable E/I balance. We found that PG cells are in fact not preferentially excited under weak OSN stimulus conditions. Nevertheless, a number of other mechanisms downstream of spiking in glomerular neurons were identified that contribute to an E/I balance that would support an intraglomerular high-pass filter.

### Input resistance and OSN input level have an equalizing effect, causing PG cells and eTCs to spike at similar levels in response to identical levels of OSN activity

Our conclusion that PG cells are not preferentially excited by weak OSN stimuli was based on two types of comparative analyses that were made between PG cells and eTCs, which are a class of glutamatergic interneurons that reside in parallel positions with respect to PG cells in the glomerular microcircuitry (**Fig. 1B**). First, in pair-cell patch-clamp recordings in PG cells and eTCs at the same glomerulus, we found that spiking in the two cell types required similar levels of electrical stimulation applied to OSNs (**Fig. 5**). Because, in these studies, the number of activated OSN axons varied as a function of the stimulation amplitude, the similar stimulation threshold indicated that individual PG cells and eTCs were activated at similar levels of OSN activity. Second, we found using calcium imaging studies in VGAT-Venus rats (Uematsu et al., 2008; Whitesell et al., 2013) that a labeled GABAergic neuron at a glomerulus was no more likely to be activated as an unlabeled neuron (presumed to be an eTC; **Fig. 6**). In the imaging analysis, weak electrical stimulation of OSNs activated ∼3 times more GABAergic cells than presumed eTCs. However, this difference appeared to reflect the fact that there are many more PG cells than eTCs (Shepherd et al., 2004; Parrish-Aungst et al., 2007) rather than activation probability differences for individual PG cells/eTCs.

That we obtained a similar conclusion about the activation probabilities of PG cells and eTCs using pair-cell electrophysiological recordings and calcium imaging was important, since each approach had distinct advantages. In the pair-cell recordings, we had information about the dendritic morphology of the test cells as well as their electrophysiological responses, which are aspects (Hayar et al, 2004b; Kiyokage et al., 2010; Gire et al., 2019) that enabled us to distinguish PG cells and eTCs not only from each other but also from other cell-types in the glomerular layer including GABAergic short-axon cells. Furthermore, based on cell morphology and physiological responses, we could confirm that the two test cells sent their dendrites to the same glomerulus and thus were activated by the same set of OSNs. On the other hand, the imaging methods enabled us to record from many more cells at once, while also providing information about whether the cells were GABAergic or non-GABAergic (based on VGAT-Venus expression). It should be noted that at least some of the VGAT+ cells analyzed with the imaging – those with larger cell bodies (see **Fig. 6Db**) – may have reflected GABAergic short-axon cells.

The observed similar activation probability for PG cells and eTCs under weak stimulus conditions was, in some sense, surprising. PG cells have long been recognized to stand out due to their very small cell bodies and dendritic arbors, along with an associated high input resistance (0.5-2 GΩ; Puopolo and Belluzzi, 1998; McQuiston and Katz, 2001; Smith and Jahr, 2002; Hayar et al., 2004b). The high input resistance in principle should make PG cells preferentially respond to weak excitatory currents (Cleland and Sethupathy, 2006; Gire and Schoppa, 2009; Fukunaga et al., 2014). Indeed, we did find that PG cells spiked in response to much lower levels of synaptic current versus eTCs (**Fig. 1**), as expected by their high input resistance. However, this advantage in terms of responsiveness was offset by the fact that, at a given level of OSN activity, PG cells received far less excitatory current than eTCs (**Fig. 3**). These results highlight the fact that, when considering a neuron’s responsiveness at a given level of OSN activity, one has to consider not only how a neuron responds to a given synaptic current level but also how much excitatory current is produced at a given level of OSN activity. There are other factors as well as, of course, that control the responsiveness of a neuron, such as its complement of voltage-gated ion channels (Liu and Shipley, 2008). Almost certainly, the small synaptic currents that we observed in PG cells was a natural consequence of their small size, i.e., the same factor that contributes to their high input resistance. Indeed, we found that PG cells had smaller OSN-EPSCs generated by a single OSN axons versus eTCs and also reduced OSN axon convergence (**Fig. 4**), both of which could naturally reflect the small dendritic arbor of PG cells.

Might our analysis have missed a subpopulation of PG cells that are especially responsive at low levels of OSN activity? This seems unlikely. While there exists PG cells with very high input resistances near 2 GΩ (Najac et al., 2015; Benito et al., 2018) – in principle making these cells especially responsive – we found that highest resistance PG cells were no more responsive than other PG cells (**Fig. 5E**). That the highest-resistance cells were no more responsive may have reflected the fact that at least some of these cells were immature and poorly integrated into the bulbar network (Benito et al., 2018). Differences in activation probabilities were also not observed when we sorted the PG cells by their mode of excitation following OSN stimulation, i.e. whether they were ONd-PG cells excited solely by OSN input or ETd-PG cells that can be activated by a polysynaptic pathway (Shao et al., 2009; **Fig. 5D**).

There are some limitations around our conclusion that individual PG cells and eTCs are activated to a similar degree by OSN activity. For example, in our analysis of PG cells, we chose to focus on fast sodium spikes as a measure of cell activation, but prior studies have suggested that, at least amongst a subset of bursting PG cells (McQuiston and Katz, 2001; Najac et al., 2015), dendritic calcium spikes drive most of GABA release (Murphy et al. 2005). How much these prior results relate to our analysis is however not clear. In their analysis of calcium spikes, Murphy and co-workers (2005) used direct depolarization of PG cells as a stimulus, which is quite distinct from the synaptic stimulus that we used. In our recordings of responses to OSN stimulation, we did not consistently observe calcium spikes in PG cells (see example traces in **Fig. 5**). An additional consideration is that, even if calcium spikes are important during a synaptic stimulus, our results still argue against the prevailing view that PG cells should be more easily excited by OSNs than excitatory neurons at glomeruli (Cleland and Sethupathy, 2006; Gire and Schoppa, 2009; Fukunaga et al., 2014). In their prior analysis, Murphy and co-workers (2005) found that calcium spikes required a stronger depolarizing stimulus to be activated versus sodium spikes; hence, by focusing on sodium spikes in our studies, we have, if anything, overestimated the degree to which a given stimulus excites PG cells to produce a GABAergic output.

### Factors that contribute to an E/I balance that support an intraglomerular high-pass filter

Our comparative analysis of the stimulus-response relationship of eTCs and PG cells argued against one potential mechanism underlying an intraglomerular high-pass filter, involving input resistance differences between these cells. However, we were able to obtain evidence for other mechanisms that could support a filter when we broadened our analysis to examine the excitatory and inhibitory currents that result secondarily from activation of eTCs and PG cells.

Our first result from this analysis was that a glomerulus does indeed display E/I balances that support an intraglomerular high-pass filter (**Fig. 1A**): Weak OSN stimuli elicited an unfavorable E/I conductance ratio (∼0.20), but ∼5-fold greater ratios were observed with strong OSN stimuli. We reported a similar shift in the E/I conductance ratios with stimulus strength in another recent paper (Gire et al., 2019), albeit using somewhat different thresholds for what constituted weak versus strong OSN stimuli (based on the size of the OSN-EPSC in the eTC). In addition, our pharmacological analyses of the eTC currents identified two factors that impact the intraglomerular E/I ratio, in opposing directions: (1) glial glutamate transporters that limit recurrent excitation; and (2) presynaptic Group II mGluRs that limit GABA release from PG cells (Zak et al., 2015). Importantly, the actions of the glial glutamate transporters were ubiquitous across OSN stimulation conditions – the transport blocker DL-TBOA enhanced the E/I ratio to a similar degree regardless of the amplitude of the OSN-EPSC – but the actions of Group II mGluRs were specific to strong OSN stimuli. Thus, glutamate transporters appear to have an overall tempering effect on excitation, contributing to lower E/I ratios across all OSN activity levels, while Group II mGluRs have an offsetting effect favoring greater E/I ratios that is selective for when OSNs are strongly activated. That the Group II mGluR-effects were limited to strong stimuli may have reflected an extrasynaptic localization for these receptors (Shigemoto et al., 1997; Schoepp, 2001). With such a positioning, the glutamate transients sufficient to activate the receptors may require activity in many glutamate-releasing neurons at once.

Our study was by no means exhaustive, and there are likely other mechanisms that could also shape the E/I balance consistent with an intraglomerular high-pass filter. For example, prior studies in eTC-MC pair-cell recordings reported a highly supralinear relationship between the number of spikes in an eTC and the extrasynaptic excitatory current (Gire et al., 2019), perhaps reflecting the dynamics of glutamate accumulation in the extrasynaptic space. A similar relationship could hold between increasing OSN activity levels and the extrasynaptic current, contributing to more favorable E/I ratios. The glomerular E/I balance would also likely be impacted by specific axonal inputs from higher-order brain centers that impinge on eTCs or PG cells (Boyd et al., 2012; Ma and Luo, 2012; Markopoulos et al., 2012; Rothermel and Wachowiak, 2014; Sanz Diez et al., 2019). Other factors contributing to the filter may also operate at a more global, cell-population level. For example, prior studies have established that there is a large discrepancy in just the size of the population of GABAergic versus non-GABAergic cells at a glomerulus, with many more GABAergic cells (Shepherd et al., 2004; Parrish-Aungst et al., 2007). That there is a greater number of GABAergic cells highlights the fact that signaling at a glomerulus may be generally weighted in favor of inhibition, and it is only under special circumstances (e.g., when Group II mGluRs are excited) that excitation can overcome inhibition.

A final point about our estimates of the E/I balance, perhaps a caveat, is that they were made in eTCs and not directly in output MCs and TCs. The choice to perform the estimates in eTCs rather than MCs was based on morphological considerations (Gire et al., 2019). While eTCs have only apical dendrites, MCs and TCs have both apical and lateral dendrites, which complicate the interpretation of the recorded currents as a measure of the intraglomerular E/I balance. Nevertheless, eTCs do mediate a major feedforward path for exciting MCs (OSN-to-eTC-to-MC; De Saint Jan et al., 2009; Najac et al., 2011; Gire et al., 2012; Vaaga and Westbrook, 2016). Hence, we expect that the E/I ratio in eTCs should greatly impact excitation of MCs.

### All PG cells receive direct OSN input

While the main purpose of our study was more functional, to establish input-output relationships for various bulbar neurons, our patch-clamp recordings also enabled us to draw one additional conclusion about basic neural connectivity. This pertained to glutamatergic connections onto the subtype of PG cells known as ETd-PG cells that receive strong polysynaptic signals from eTCs (Shao et al., 2009). While prior studies have suggested that ETd-PG cells receive at most “weak” direct OSN input (Shao et al., 2009; Najac et al., 2015), our recordings suggest that direct OSN inputs onto ETd-PG cells are quite robust (**Fig. 2**). At least part of the explanation for the somewhat differing conclusions is how input strength was defined. In the study by Shao and co-workers (2009), for example, the direct pathway for exciting PG cells was defined to be weak since a stronger OSN stimulus was required for its activation versus the polysynaptic pathway. On the other hand, we performed a side-by-side comparison between ETd-PG cells and ONd-PG cells, which are the subtype of PG cells that are distinguished by the fact that they only receive monosynaptic input. At similar, moderate OSN stimulation intensities, the OSN-EPSCs in ETd-PG cells and ONd-PG cells were both large (generally >200 pA) and indistinguishable from each other in size. Notably, our analysis of the convergence properties of OSNs onto PG cells versus eTCs (**Fig. 4**) also help explain the observation of Shao and co-workers (which we also made) that OSN-EPSCs in ETd-PG cells require stronger OSN stimuli to be generated than polysynaptic signals. Because eTCs receive input from ∼3-4-fold more OSN axons than PG cells, fewer OSN axons may need to be activated to activate the OSN-to-eTC-to-PG cell pathway versus the direct OSN-to-PG cell pathway. Overall, we conclude that, while the polysynaptic pathway may obscure the direct OSN pathway during analysis of electrophysiological recordings, ETd-PG cells in fact receive a large number of direct OSN inputs.

## Acknowledgements

The authors thank members of the Schoppa lab for helpful discussions.

## Additional Information

### Competing Interests

None of the authors have any conflicts of interest associated with the studies described in this manuscript.

### Funding

This work was supported by NIH grants R01DC006640 (NES) and F31DC009369 (JDZ).

### Author Contributions

All experiments were conducted in the lab of Dr. Nathan Schoppa at the University of Colorado Anschutz Medical Campus.

JDZ: Conception and design of study; acquisition, analysis, and interpretation of data; drafting and editing of manuscript.

NES: Conception and design of study; analysis and interpretation of data; drafting and editing of manuscript.

Both authors approved the final version of the manuscript and agree to be accountable for all aspects of the work in ensuring that questions related to the accuracy or integrity of any part of the work are appropriately investigated and resolved. All persons designated as authors qualify for authorship, and all those who qualify for authorship are listed.

### Contact for Research Governance at UCAMC

Dr. Thomas Flaig, Vice Chancellor for Research

### UCAMC Institutional Animal Care and Use Committee protocol number 00144

## Notes

### Competing Interest Statement

The authors have declared no competing interest.

